# Triggered and Spontaneous Dormancy in Bacteria During Feast-Famine Cycles with Stochastic Antibiotic Application

**DOI:** 10.1101/2025.02.26.640324

**Authors:** Silja Borring Låstad, Namiko Mitarai

## Abstract

Bacteria can enter dormancy triggered by stress, such as starvation. When stress is removed, a large part of the population will exhibit some lag time before regrowth. It has been observed that even under stress-free conditions that allow for exponential growth, a small subpopulation can spontaneously enter dormancy temporarily. The dormant population often survives antibiotic application because many types of antibiotics target the cell growth and division process. If bacteria are in an environment where antibiotics are *sometimes* applied, the population can evolve to adjust their dormancy frequency to better survive antibiotics without losing too much of the population growth. Here, we consider the situation in which antibiotics are applied stochastically during repeated feast-famine cycles. We analyse the best strategy for long-term growth when the bacteria are allowed to tune both the lag time at the start of the feast period and the spontaneous dormancy in the feast period. We show that spontaneous dormancy can provide an advantage only when the antibiotic application and the start of the feast period are decoupled. When the triggered dormant and spontaneous dormant states are treated as different states and the antibiotic addition time is fixed, the optimal strategy is either triggered or spontaneous dormancy. Exhibiting both types of dormancy is optimal only when there is a certain level of stochastic fluctuation in the antibiotic application timing.

## I. INTRODUCTION

It is well known that dormant or slow-growing bacteria are more tolerant to multiple antibiotics [1–3]. This is because most antibiotics target cell growth and division processes, such as cell wall synthesis and DNA replication. Therefore, the efficacy of antibiotic killing depends on the bacterial growth phase. Exponentially growing bacteria are the most sensitive to antibiotics. Less killing is observed when bacteria are in the stationary phase (e.g., due to a lack of nutrients) or in the lag phase (e.g., the dormant state just after adding nutrients to the stationary phase cells) [4, 5]. In addition, even in the exponential growth phase, a small fraction of cells often survive the application of antibiotics [6, 7]. In some cases, those cells are found to be in a dormant, non- or slowly growing state [6].

The subpopulation of bacteria that survives the antibiotic application due to their phenotypic difference is often termed persisters (for a more precise definition of persisters, refer to ref. [1]). Persisters triggered by non-lethal stresses are called triggered persisters, while the persisters found during the steady-state exponential growth phase are called spontaneous persisters. In general, spontaneous persisters are much less frequent than their triggered counterparts [7, 8]. Evolution experiments have shown that persistence frequency is an evolvable trait that adapts to both the duration and frequency of the antibiotic application [4, 9]. Furthermore, persistence can promote antibiotic resistance because survival by persistence can allow bacteria to mutate to gain antibiotic resistance [10]. Hence, finding an antibiotic administration strategy that minimizes the evolution of persistence is crucial.

The dormancy-based persistence has been analysed from the game-theoretical point of view [11]. Spontaneous persistence during a continuous exponential growth phase can be interpreted as a bet-hedging behaviour, i.e., investing a small fraction of the population in dormancy to survive unpredictable lethal stress [9, 12, 13]. The optimal strategy depends on the duration of the stress; e.g., when the antibiotic is applied periodically, the stochastic switching between dormancy and growth, which provide spontaneous persistence, is beneficial only if the stress duration is long enough [14]. For triggered persistence, a periodic feast-famine cycle with deterministic application of the antibiotics were thoroughly analysed by using an individual based model [15]. Stochastic application of antibiotics synchronized with nutrient additions has also been analysed [16], and it was found that investing in a subpopulation with a prolonged lag time is expected to be the best strategy when the antibiotic application is frequent and lethal. However, as far as we know, the best strategy where both spontaneous and triggered persistence are allowed in the feast-famine cycle has not been analysed before.

In this paper, we extend the previous work of triggered persistence [16] to ask what is the best strategy when the population can exhibit both triggered and spontaneous persistence. We focus on the case where the population experiences repeated feast-famine cycles, but the timing of the addition of antibiotics and the addition of nutrients is not necessarily synchronized. In such a case, it is not apparent how to combine the two types of persistence to achieve the highest long-term population growth. We first introduce a two-state model, where triggered dormancy and spontaneous dormancy are treated as the same state. The model predicts three phases of optimal strategies depending on the timing, frequency, and duration of the antibiotic application. Here, either (i) no dormancy, (ii) only triggered dormancy, or (iii) both triggered and spontaneous dormancy are optimal. We then extend the model to a three-state model, where the triggered dormancy (lag-time) and spontaneous dormancy are treated as separate states. This model also predicts three phases for the optimal strategy when antibiotic addition timing is fixed, which are (i) no dormancy, (ii) only triggered dormancy, or (iii) only spontaneous dormancy. We observe the coexistence of triggered and spontaneous dormancy in the three-state model only when the time interval between the nutrient addition and the addition of antibiotics fluctuates sufficiently.

## II. TWO-STATE MODEL WITH TRIGGERED AND SPONTANEOUS DORMANCY

### A. Two-state model

We consider feast-famine cycles with stochastic application of antibiotics. When nutrients are available, the bacteria can be either in the dormant state or in the growth state. Cells in the dormant state do not divide but will not die if an antibiotic is applied, whereas cells in the growth state will divide when the environment is antibiotic-free but die at a constant rate if an antibiotic is applied. We first analyse the model where the triggered dormancy (lag phase) and the spontaneous dormancy are treated as the same state for simplicity, and we call it the two-state model. We later extend the model to distinguish the two dormant states.

The feast period begins when a fixed amount of nutrients *S*_0_ is added to the system. Then, the non-dormant bacteria grow, consume all the nutrients, and re-enter the dormant, stationary phase. The bacteria are then diluted *f* -fold before the next addition of nutrients. Hence, when the nutrient is added, the entire population is in the stationary dormant phase. This defines one feast-famine cycle. Note that this setup is motivated by the previous experiment [4].

We consider multiple (*n*) species competing for nutrients to find the optimal strategy in this setup. Denoting the available nutrients at time *t* as *S*(*t*) and the population size of the dormant (growth) state of species *i* as *d*_*i*_(*t*) (*g*_*i*_(*t*)), our model equation for *S*(*t*) *>* 0 is given by

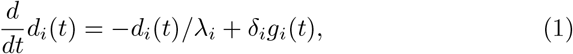

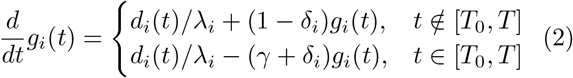

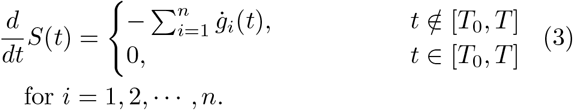

Here, we measure the nutrients in the unit that a bacterium consumes to divide once. We assume that the antibiotic is applied at time *T*_0_, lasting for a duration *T*_*ab*_ ≡ *T* − *T*_0_, with probability *p*, and otherwise set *T*_*ab*_ = 0 to represent the stochastic application of the antibiotic. The parameter 1*/λ*_*i*_ characterizes the exit rate from the dormant state, and *δ*_*i*_ characterizes the rate of spontaneous entry to dormancy when the nutrient is available. We set the growth rate of the bacteria to be the same for all species and set it to one. This defines the unit time in the model. The death rate in the presence of antibiotics is parametrised by *γ*.

As the initial condition, all populations are assumed to be in the stationary phase, i.e., *d*_*i*_(0) = *d*_0_ *>* 0 and *g*_*i*_(0) = 0. Each round of the feast period starts with *S*(0) = *S*_0_. The nutrients are consumed according to eq. (3), which ensures competition between the species. Note that in this setup, bacteria consume nutrients both when they grow and when they undergo phenotypic switching to the growth state, though the effect of the latter is negligible (see Supplementary Sec. S1). The feast phase ends when *S*(*t*) reaches zero and all variables stop changing. In practice, we simulate the system for a long enough time *t*_*f*_ so that *S*(*t*_*f*_) = 0 is reached. We then calculate the total population for each species in the final stage of the cycle *p*_*i*_(*t*_*f*_) = *d*_*i*_(*t*_*f*_) + *g*_*i*_(*t*_*f*_). Before starting the next round, we put a fraction *f* of the surviving population from the previous round *p*_*i*_(*t*_*f*_) in the dormant state, i.e., *d*_*i*_(0) = *f · p*_*i*_(*t*_*f*_) and *g*_*i*_(0) = 0. We then set *S*(0) = *S*_0_ and start the next feast round. We repeat this cycle multiple times to select the best strategy.

In the following, we set the death rate *γ* = 1; that is, we assume it to be comparable to the growth rate. We set *S*_0_ = 10^9^ cells/ml, *d*_0_ = 10^3^ cells/ml and the dilution fraction *f* = 10^−6^ unless otherwise noted.

### B. Two-state model predicts three phases of optimal strategies

If there is only one species in the environment, dormancy does not bring any disadvantage because any growing population will eventually increase by *S*_0_ every round. This changes when there is competition: Fig. 1 shows an example trajectory of competition between two species under a stochastic antibiotic application with frequency *p* = 0.4. Here, species *p*_1_ has a relatively long lag time and no spontaneous dormancy, while species *p*_2_ has a short lag time and a finite rate of entering spontaneous dormancy. We see that species *p*_1_ is dominant when there is an antibiotic in the environment (the first and fourth cycle). The relative population of species *p*_2_ increases when there is no antibiotic, though not enough to “win” the competition. After many repetitions, species *p*_1_, with longer lag time and no spontaneous persistence, will dominate in this example of relatively rare antibiotic application.

**FIG. 1.**
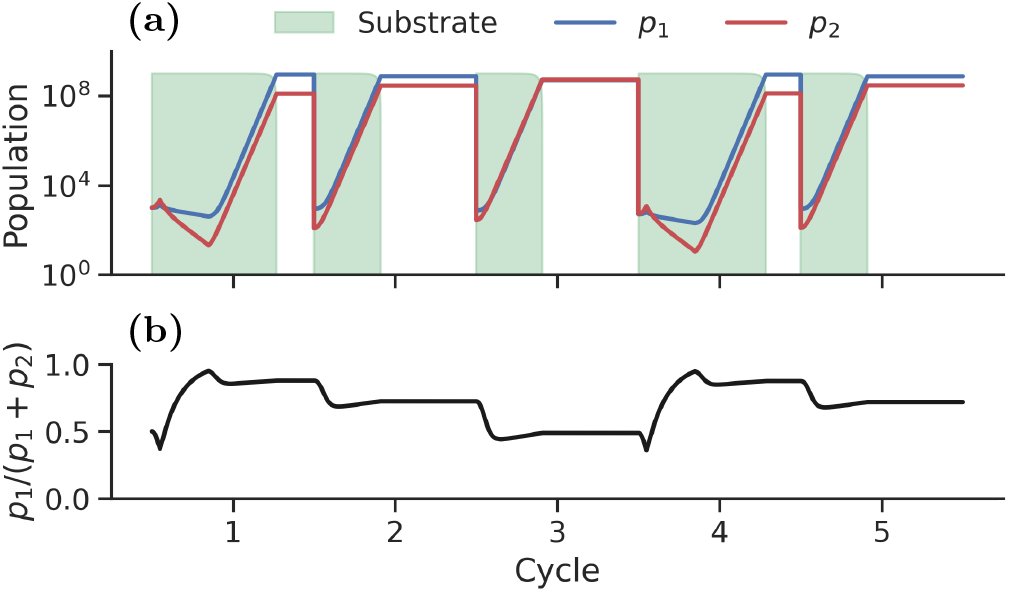
An example trajectory of five feast-famine cycles with two species competition. Here, *p* = 0.4, *T*0 = 2, and *T*_*ab*_ = 12. (a) The blue line represents species *p*_1_ with *λ*_1_ = *T* and *δ*_1_ = 0 (no spontaneous dormancy), while the red line shows species *p*_2_ with *λ*_2_ = 3 and *δ*_2_ = 0.04. The shaded light green area shows the nutrient level. (b) Intra-population frequency dynamics plotted as the population of species *p*_1_ over the total population of both species *p*_1_ + *p*_2_.

However, it is difficult to perform analytical calculations of the optimal strategy in an explicit competition of two or more species. Here, we first present an analysis considering the single species “optimal” strategy: We consider a single species in our setup and find the persistence parameters that minimize the average time it takes the species to consume all nutrients per round ⟨*T*_*s*_⟩. The average is taken over multiple rounds of feast-famine cycles.

When there is only one species, it is possible to solve the model equations (1)-(3) analytically to obtain the time taken to consume all nutrients with and without antibiotic application, respectively (Appendix A). Since the system experiences a round of antibiotics with probability *p*, we take the average of these times with respect to the weights *p* and 1 − *p*. The final expression is given by (we omit the subscript for species identity)

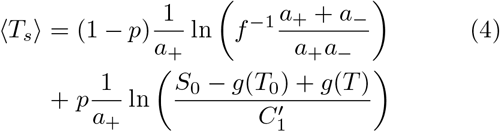

where

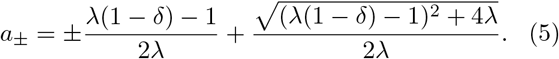

*g*(*T*_0_), *g*(*T*) and 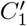 are also functions of the species parameters *λ, δ*, and *γ*, and their functional forms are given in Appendix A.

We maximize the effective single-species fitness

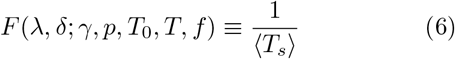

numerically to predict the optimal strategy. More specifically, we find a set of values (*λ, δ*) = (*λ*^⋆^, *δ*^⋆^) that yields the largest *F* for a given set of environmental parameters *γ*(= 1), *p, T*_0_, *T* and *f* (= 10^−6^). The obtained optimal parameters are presented in Fig. 2, with the corresponding *F* in Supplementary Fig. S3. In Fig. 2a-b, we fixed the duration of the antibiotic application *T*_*ab*_ = 10 and varied the probability *p* and the starting time *T*_0_ of the antibiotic application after nutrient addition to find the optimal (a)*λ*^⋆^ and (b)*δ*^⋆^. Interestingly, we observe three phases: When *p* is small, *λ*^⋆^ = 0 and *δ*^⋆^ = 0, i.e., neither triggered nor spontaneous dormancy gives enough advantage to compensate for the loss of growth associated with dormancy. For a given *T*_0_, there is a critical value of *p* at which it becomes beneficial to enter dormancy. When *T*_0_ = 0, that is, when the antibiotics are added at the same time as the nutrients, it is best to have only triggered persistence, with *λ*^⋆^ ∼ *pT >* 0 and *δ*^⋆^ = 0 for large *p*. It should be noted that similar behaviour was also observed in the previous analysis without spontaneous dormancy and *T*_0_ = 0 [16]. When *T*_0_ is large enough, it becomes beneficial to wake up earlier from starvation by having a smaller, but still positive, *λ*^⋆^ to grow for *t < T*_0_. Here, it is also beneficial to allow the bacteria to spontaneously re-enter dormancy by setting *δ*^⋆^ *>* 0 to hedge the bet for large antibiotic application probability *p*.

**FIG. 2.**
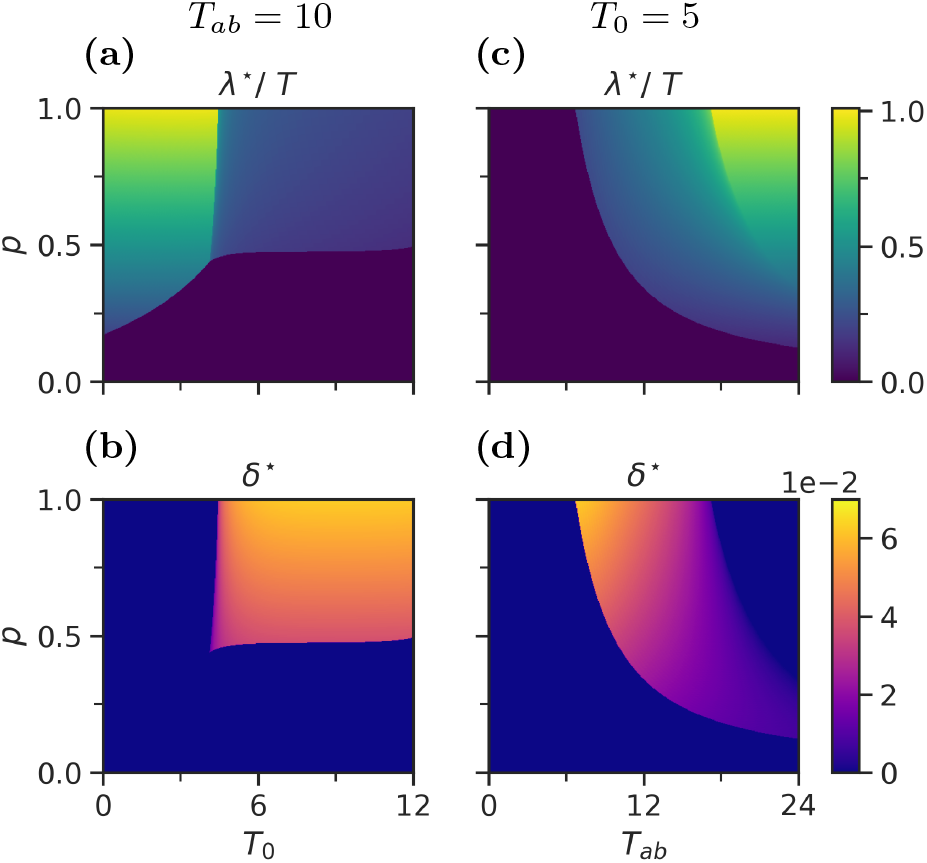
The optimal persistence parameters for the two-state model for varying *p* with *T*_*ab*_ = 10 and varying *T*_0_ (a-b) and with *T*_0_ = 5 and varying *T*_*ab*_ (c-d). (a,c) The optimal lag time (*λ*^⋆^) normalized by *T*, and (b,d) the rate to enter spontaneous dormancy (*δ*^⋆^).

We also performed a similar analysis with fixed *T*_0_ = 5 and varying the duration of the antibiotic application *T*_*ab*_ (Fig. 2c-d). Again, we see the three phases: When *T*_*ab*_ is small enough, *λ*^⋆^ = *δ*^⋆^ = 0 is the best strategy. When *T*_*ab*_ and *p* are very large, the best strategy is to have triggered persistence only, with *λ*^⋆^ ∼ *pT >* 0 and *δ*^⋆^ = 0. For the middle value of *T*_*ab*_ (including *T*_*ab*_ = 10) and sufficiently high *p*, we observe the best strategy to be positive *δ*^⋆^ and *λ*^⋆^. We tested whether the best strategy obtained based on the single species fitness eq. (6) matches the optimal strategy obtained by direct competition of two different strategies, i.e., two sets of (*λ, δ*) values. For a given set of environmental parameters, we chose the optimal set of single species parameters (*λ*^⋆^, *δ*^⋆^) and scanned the competitor parameter space as follows: *λ* ∈ [0.01, *T*] with a linear interval of *T/*100, and *δ* ∈ [0, 0.1] with a linear interval of 0.001. This yielded 100 *×* 100 types of competitors that the optimal strategy (*λ*^⋆^, *δ*^⋆^) competed against one by one. The competition was carried out over 10^4^ feast-famine cycles, where we considered the cycle average of how much nutrients were converted to the biomass of a given species as a measure of competition fitness. The optimal competitor is then the strategy that consumes the largest amount of nutrients, where this fraction approached 1 (Supplementary Sec. S4).

The best competitor strategies are compared to the result based on eq. (6) in Fig. 3. We see that the agreement is reasonably good and we observe the same three phases with discontinuous and continuous transitions depending on where we cross the boundary. In Fig. 3a-b (*T*_*ab*_ = 10), the competition and single species optimal parameters overlap almost perfectly, but the optimal rate to enter spontaneous dormancy is higher when there is competition. In Fig. 3c-d (*T*_0_ = 5), there is also a significant shift in the phase boundary between the mixed dormancy strategy and the purely triggered strategy, in addition to the increase in *δ*^⋆^. Competition has qualitatively the same effect on the optimal persistence parameters as increasing the dilution fraction *f* (details in Supplementary Sec. S5), which is reasonable since both effectively shorten the consumption time. These results were also confirmed in a competition simulation of mutation populations (See Supplementary Sec. S6). In this case, where several species compete for the nutrients, the best competitor strategies mostly dominate, though often with large fluctuations around the optimal values.

**FIG. 3.**
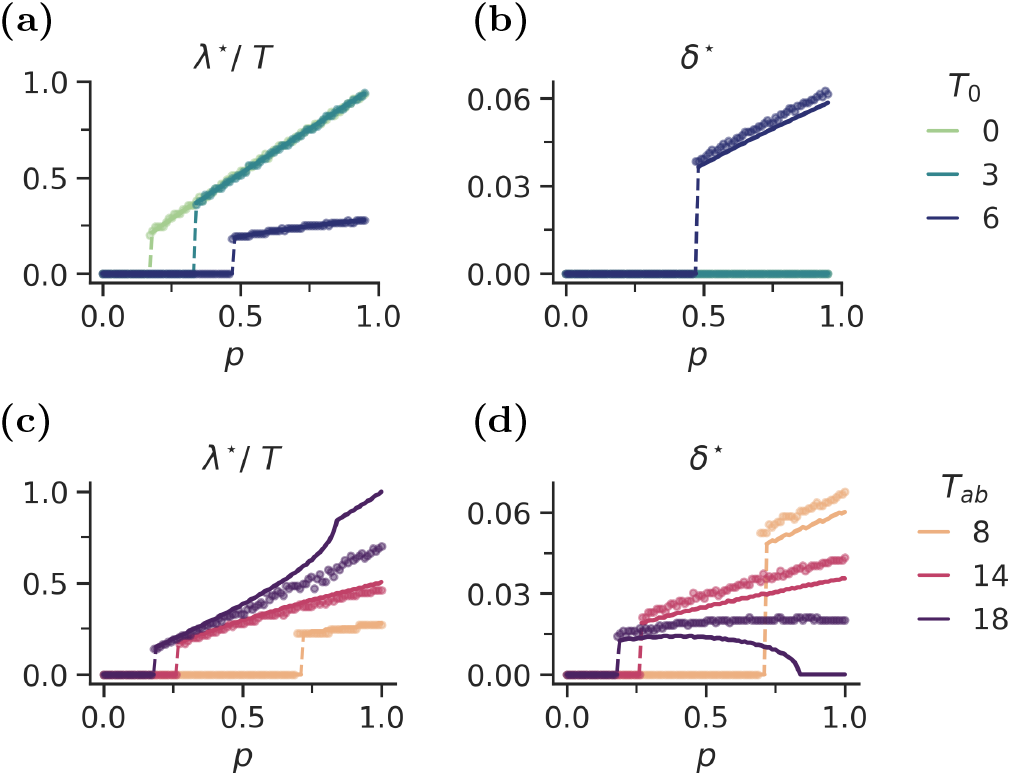
Optimal lag time and optimal rate of spontaneous persistence obtained as a function of the antibiotic application probability *p* by direct competition of strategies. (a) optimal lag time *λ*^⋆^ and (b) optimal rate of spontaneous dormancy *δ*^⋆^ for *T*_*ab*_ = 10. (c) optimal lag time *λ*^⋆^ and (d) optimal rate of spontaneous dormancy *δ*^⋆^ for *T*_0_ = 5. The circles denote the competition optima, and the solid lines denote the single species optima from *F* (eq. 6).

## III. THREE-STATE MODEL: TRIGGERED AND SPONTANEOUS DORMANCY AS SEPARATE STATES

### A. Three-state model

So far, we have treated triggered and spontaneous dormancy as the same state. However, given the multiple possible molecular mechanisms behind dormancy and their diverse physiological states [17–25], it is likely that the lag state after starvation and the spontaneous dormancy state are different states, though they could be related. Next, we therefore analyse a model where we distinguish the two dormant states and allow each to have a different wake-up rate. We denote the growing population as *g*_*i*_(*t*), the triggered dormant population as *d*_*i*_(*t*), and the spontaneously dormant population as *r*_*i*_(*t*). The resulting model equations are

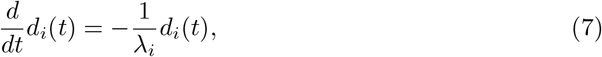

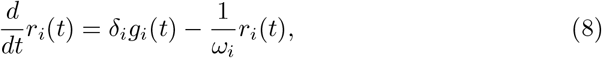

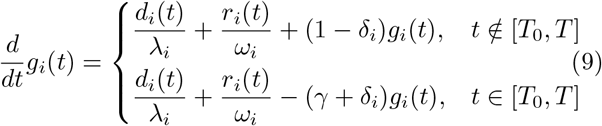

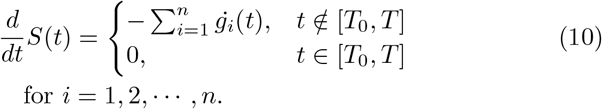

Here, the wake-up rate from spontaneous dormancy is set by the parameter *ω*_*i*_. The rest of the model’s setup, such as the nutrient addition and the antibiotic application, is the same as in the two-state model.

An example trajectory of competition between two species in the three-state model is shown in Fig. 4. species *p*_1_ has long lag time and no spontaneous dormancy, whereas species *p*_2_ has only spontaneous dormancy. The antibiotic parameters are the same as in Fig. 1, but in contrast to the two-state model we see that species *p*_2_ is dominant in the three-state example trajectory.

**FIG. 4.**
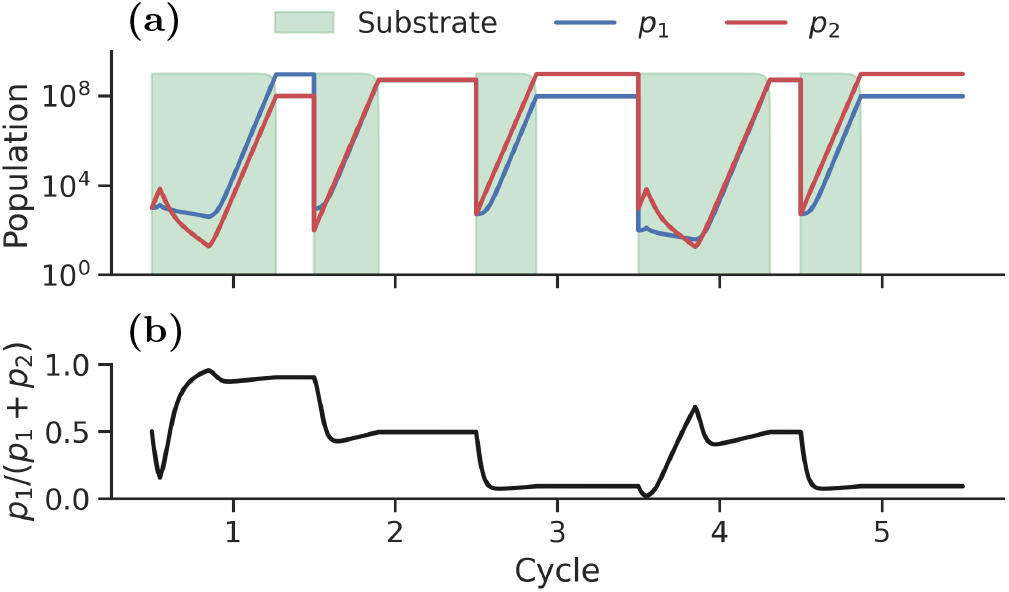
An example trajectory of five feast-famine cycles with two species competition. Here, *p* = 0.4, *T*_0_ = 2, and *T*_*ab*_ = 12. (a) The blue line represents species *p*_1_ with *λ*_1_ = *T, ω*_1_ = 0.01 and *δ*_1_ = 0 (no spontaneous dormancy), while the red line shows species *p*_2_ with *λ*_2_ = 0.01, *ω*_2_ = 3, and with *δ*_2_ = 0.04. The shaded light green area shows the nutrient level. (b) Intra-population frequency dynamics plotted as the population of species *p*_1_ over the total population of both species *p*_1_ + *p*_2_. Notice that species *p*_2_ here appears to have an advantage, which is the opposite of what we observe with the same antibiotic setup in Fig. 1.

### B. The best strategy is to choose either triggered or spontaneous persistence in three state model

The analytical estimate of the optimal parameter set for the three-state model is shown in Fig. 5, with the corresponding *F* in Supplementary Fig. S10. Similarly to the two-state model, the optimal set of parameters (*λ*^⋆^, *ω*^⋆^, *δ*^⋆^) was estimated as the set that minimizes 1*/*⟨*T*_*s*_⟩, in the three-state model defined as

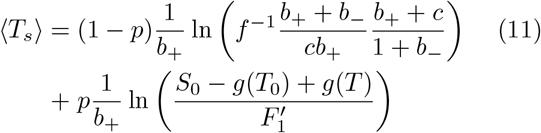

where

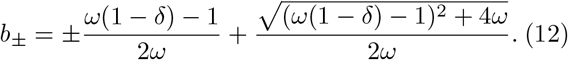

where *g*(*T*_0_), *g*(*T*) and 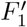 are given in Appendix B.

First, we analysed the three-state model with *T*_0_ = 0 (Fig. 5a-c), i.e., when the antibiotic is added at the same time as the nutrients. When we look at the optimal lag time *λ*^⋆^, it shows a first-order transition, which is qualitatively similar to the triggered persistence-only model [16]. However, for a sufficiently long antibiotic application time *T*_*ab*_, we see that a low but positive spontaneous persistence rate *δ*^⋆^ becomes favourable, with wakeup time *ω*^⋆^ comparable to *T*. In other words, if *T*_*ab*_ is very long, it pays to have a small fraction in the spontaneous persisted state, even if the probability of antibiotic application *p* is very low.

**FIG. 5.**
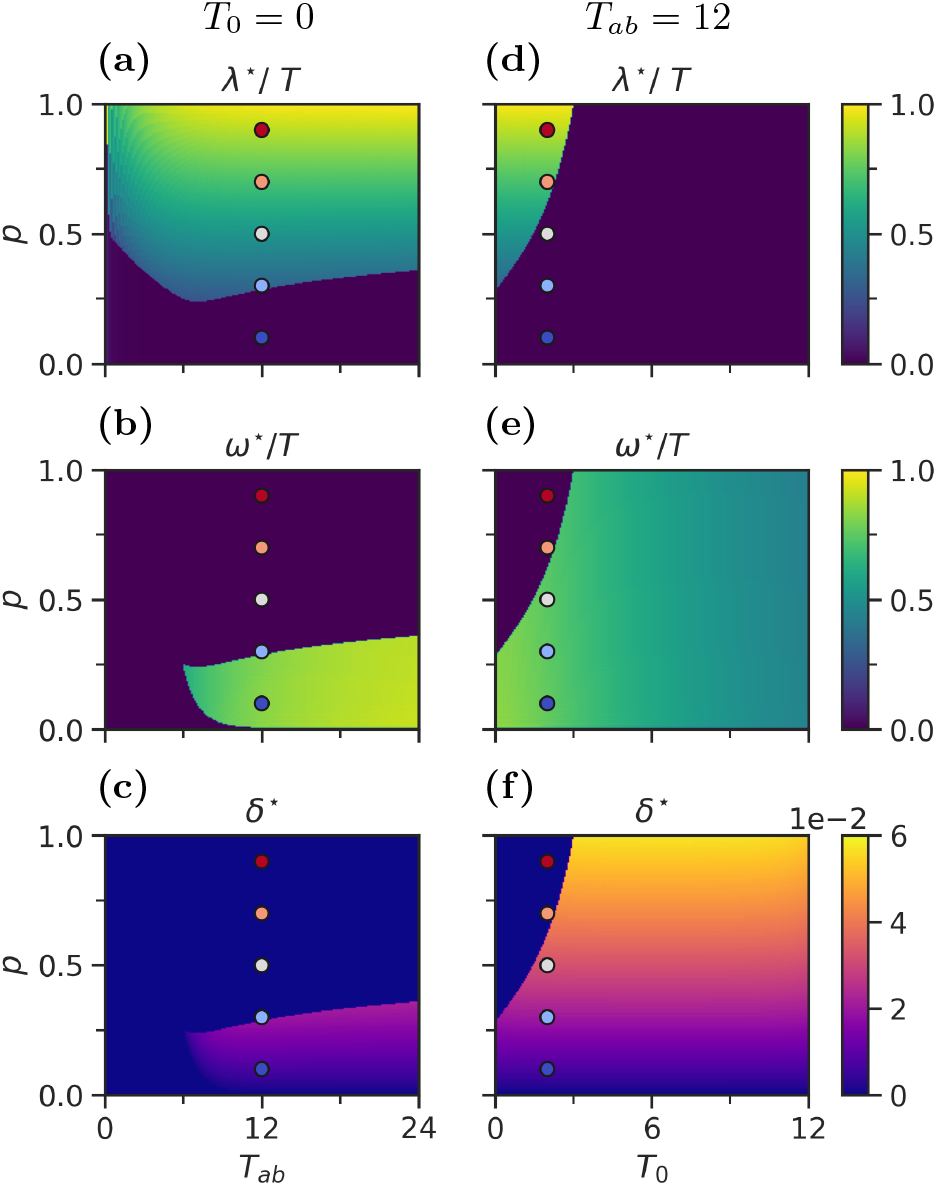
The optimal parameters for three-state model for varying *p* with *T*_0_ = 0 and varying *T*_*ab*_ (a-c) and with *T*_*ab*_ = 12 and varying *T*_0_ (d-f). (a,d) The lag time (*λ*^⋆^) normalized by *T*, (b,e) the time spent in spontaneous dormancy (*ω*^⋆^) normalized by *T*, and (c,f) the rate to enter the spontaneous dormancy (*δ*^⋆^). The coloured circles indicate the parameters used in the competition simulation in Fig. 6.

Next, we explored the case where *T*_*ab*_ = 12, with varying *T*_0_ and *p* (Fig. 5d-f). We again see the separation of the parameter range; *λ*^⋆^ ∼ *pT >* 0 for small *T*_0_ and large *p*, while in that region *δ* = 0, i.e., there is only triggered persistence. For the other region, *λ*^⋆^ = 0, and *δ*^⋆^ is small but finite, with *ω*^⋆^ ≈ 0.5*T*. This is somewhat similar to the two-state model case (Fig. 2), where there was a clear separation of the *δ*^⋆^ = 0 region and the *δ*^⋆^ *>* 0 region. However, in the two-state model *λ*^⋆^ *>* 0 when *δ*^⋆^ *>* 0, since it also controlled the wake-up rate of the spontaneous dormant state. In the three-state model, we see that the best strategy is either to have triggered or spontaneous persistence.

We speculated whether the phase of the mixed persistence strategy (*λ*^⋆^ *>* 0 and *δ*^⋆^ *>* 0) could also represent an optimal strategy in the three-state model if we let *T*_0_ fluctuate. In this case, the bacteria are exposed to both the situation where triggered persistence is more advantageous and *vise versa*. The result is summarized in supplementary Fig. S11 and S12. We, in fact, observed that the mixed strategy is optimal if the fluctuation amplitude is sufficiently large and the low *T*_0_ (which favours triggered dormancy) is more frequent than the high *T*_0_ (spontaneous dormancy).

We then compared the phase diagram in Fig. 5 to a competition simulation of mutating populations. We let a small subpopulation with slightly altered parameters appear and let them compete to find the optimal parameter set. This “mutation” process was implemented by a low mutation rate *ε* = 10^−3^ upon the population growth as follows: We let a population to have 3 indices (*i, j, k*), which has a parameter set

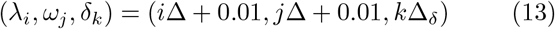

with Δ = 0.5 and Δ_*δ*_ = 0.005. We then assume that upon mutation, the parameters can evolve to the nearest neighbour values for a set of integers (*i, j, k*). Then, in the absence of antibiotics, the growing part of the population (*i, j, k*) obeys

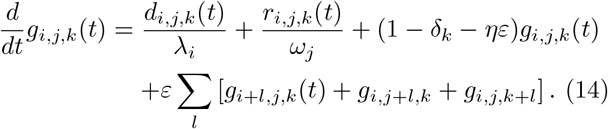

Here, *l* ∈ {−1, 1} if both neighbour values exist. We limit the minimum values of the parameters to (0.01, 0.01, 0). Hence, the range of *l* is limited when one or more indices are 0. In addition, we put an upper bound on the parameters at (*T, T*, 0.1) and limit the range for species (*λ*_*i*_, *ω*_1_, *δ*_1_) because mutations to *ω*_0_ and *δ*_0_ always occur simultaneously (i.e., species *g*_*i*,1,0_ and *g*_*i*,0,1_ do not exist). *η* then reflects the number of neighbour states to which a parameter set can mutate, ranging between 2 and 6. We let the system run over 10^4^ feast famine cycles, starting with only species (0, 0, 0) and observed how the average parameters evolved.

The results are shown in Fig. 6. The panel (a-c) represents the case of synchronized nutrients and antibiotics, with *T*_0_ = 0 and *T*_*ab*_ = 12. We observe that the result is consistent with Fig. 5a-c, with only triggered persistence for high frequencies of antibiotics (*p* ≥ 0.5). In Fig. 6c, we observe ⟨*δ*⟩ *> δ*^⋆^ = 0, which is expected since *δ* = 0 is the lower limit of the mutation range. The fluctuations around the optimal values decrease with increasing *p*, highlighting that the optimal strategy becomes more dominant with high *p*. At *p* = 0.3, we are very close to the phase boundary, and we observe large fluctuations in all parameters. Both ⟨*λ*⟩ and ⟨*ω*⟩ are centred around the expected values for triggered persistence, but the variables sometimes ‘switch’ to spontaneous persistence for a few cycles. However, ⟨*δ*⟩ does not simply fluctuate between its triggered and spontaneous optima. When switching from triggered to spontaneous persistence, ⟨*δ*⟩ increases first before relaxing towards its minimum. At *p* = 0.1, we observe mainly spontaneous persistence, but here, we also observe an increase in ⟨*δ*⟩ when the fluctuations in the other parameters are approaching triggered persistence. This indicates that a high rate of entry to spontaneous dormancy can provide increased protection during the vulnerable transition between the two optima.

**FIG. 6.**
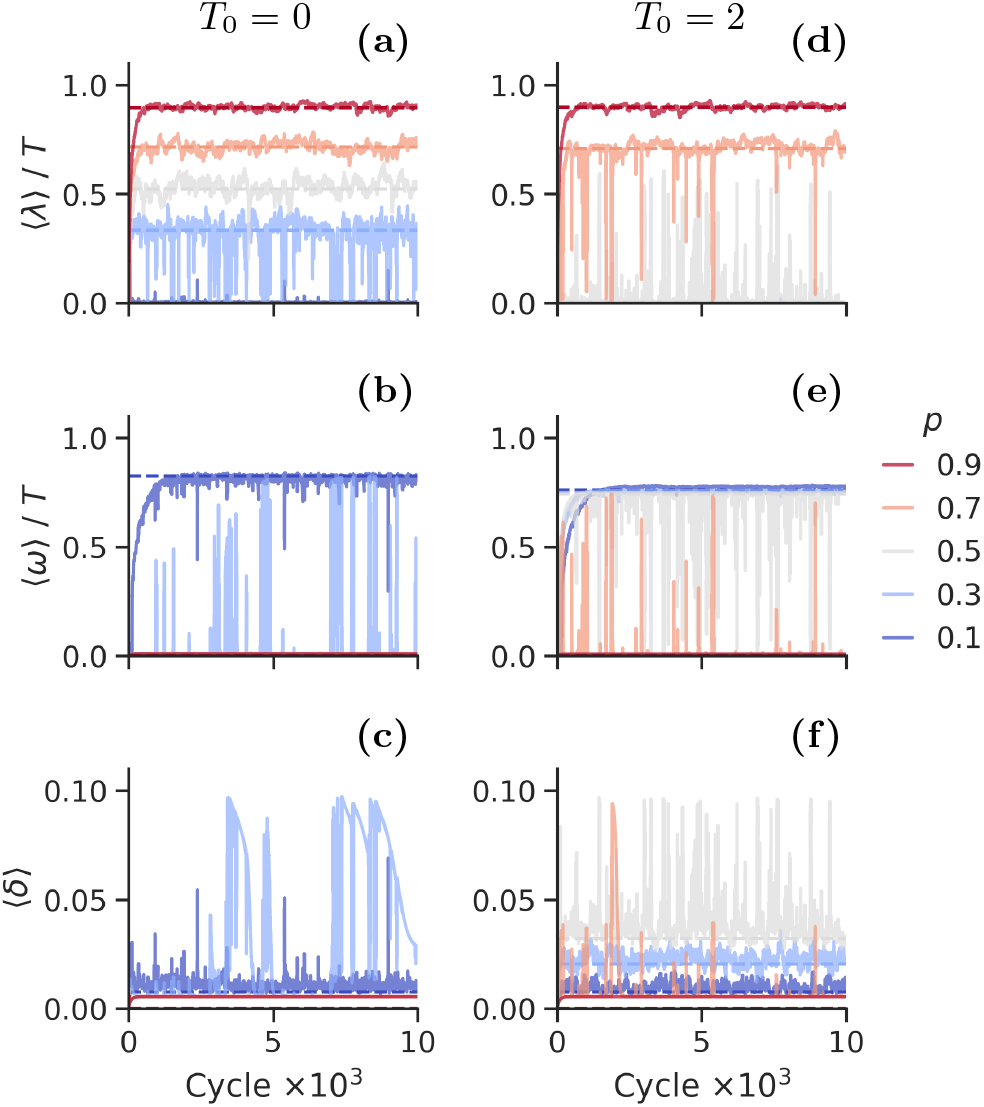
Evolution of average persistence strategy for *T*_0_ = 0 and *T*_*ab*_ = 12 (a-c) and *T*_0_ = 2 and *T*_*ab*_ = 12 (d-e) with varying *p* and the mutation rate *ε* = 10^−3^. (a,d) The average lag time (⟨*λ*⟩) normalized by *T*, (b,e) the time spent spontaneous dormancy (⟨*ω*⟩) normalized by *T*, and (c,f) the rate to enter the spontaneous dormancy (⟨*δ*⟩). Different lines correspond to different values of *p*, as labelled to the figure’s right. The set of antibiotic parameters explored are highlighted as coloured circles in Fig. 5.

In Fig. 6d-f, the application of antibiotics is desynchronized from the nutrient addition, with *T*_0_ = 2 and *T*_*ab*_ = 12. The result is again consistent with the phase diagram in Fig. 5, where spontaneous persistence is preferred for *p* ≤ 0.3 and triggered persistence is preferred for *p* = 0.9, while strong fluctuations are observed around the phase boundary *p* = 0.5. For *p* = 0.7, the system tends to have a high lag time - low spontaneous dormancy state, but we still observe some fluctuations. In Supplementary Sec. S9 we explore the case of *ε* = 10^−2^, which is mainly found to increase the frequency and amplitude of fluctuations around the single species optimal values.

The large fluctuations of the parameters in the evolution simulation are consistent with the corresponding fitness values *F*, which are shown in Supplementary Fig. S14. We see that the fitness difference between the triggered dormancy and the spontaneous dormancy strategies is rather small for the parameters for which we observe frequent switching in Fig. 6. Furthermore, especially in the case of *T*_0_ = 2 with *p* = 0.5 and *p* = 0.7, the fitness landscape is very flat (Supplementary Figs. S15-S16), consistent with the large fluctuations observed.

## IV. SUMMARY AND DISCUSSION

We have analysed the optimal persistence strategy in feast-famine cycles with stochastic application of antibiotics. We extended previous work [16] to add the possibility of entering dormancy spontaneously, and we also studied the case where the application of antibiotics is delayed compared to the addition of nutrients. In the two-state model where starvation-triggered dormancy and spontaneous dormancy are treated as the same state and, therefore, have the same wake-up rate (1*/λ*), we observed that the finite rate *δ* to spontaneously re-enter dormancy becomes optimal for a sufficiently long delay in the antibiotic application *T*_0_, with appropriate duration *T*_*ab*_ and frequency *p* of antibiotic application.

When triggered and spontaneous dormancy are treated as separate states, each having a different wake-up rate, we found that the best strategy was to have no dormancy (*λ*^⋆^ = 0 and *δ*^⋆^ = 0; low enough antibiotics severity), to have triggered dormancy only (*λ*^⋆^ ∼ *pT >* 0 and *δ*^⋆^ = 0; not too long *T*_0_), or to have spontaneous dormancy only (*λ*^⋆^ = 0, *δ*^⋆^ *>* 0 and *ω*^⋆^ *>* 0; long enough *T*_0_ and/or long enough *T*_*ab*_ with small *p*). The separation of the best strategy can be understood as follows: the advantage of having *λ < T*_0_ (waking up before antibiotics are applied) comes from the growth for *t < T*_0_; therefore, this advantage is the largest when the species wakes up immediately, i.e., *λ* → 0. Then, spontaneous persistence offers some advantages for *t > T*_0_. On the other hand, if the antibiotic treatment is too severe to favour *λ* → 0, it is better to sleep through the entire antibiotic treatment, in which case spontaneous persistence offers no further advantage. The mixed strategy (*λ*^⋆^ *>* 0 and *δ*^⋆^ *>* 0) was observed only when *T*_0_ fluctuates stochastically.

We have also confirmed that the optimal strategy obtained by minimizing the average time to consume the nutrient as a single species, ⟨*T*_*s*_⟩, is mostly consistent with the evolution simulation of the optimal strategy. Here, it is important to point out the lack of extinctions in our model. If we include extinctions by assuming a lower limit of the population size, it becomes crucial for the population not to drop below the extinction threshold; hence, it might be necessary to avoid the risk of significant population decrease during rare but severe antibiotic applications, at the cost of a higher growth rate.

It is also worth noting that in the current model, spontaneous dormancy is automatically a bet-hedging strategy since only a small subpopulation is in the dormant state when the majority is growing, while the triggered dormancy is modelled as the common strategy for the entire population. In [16], a version of the model was studied in which two subpopulations can have different values of *λ*, and it was found that the optimal strategy can be for a fraction of the population to have *λ* = 0 and the rest to have a finite *λ*, which allows cells to bet-hedge also in the triggered persistence. Analysing such an option is an interesting future project.

Another interesting extension of the current model is the effect of antibiotics on dormancy. The current model with antibiotic-independent switching rates implicitly assumes that the applied antibiotic is at a lethal level and hence, the cells do not have time to adapt to it. However, there are situations where antibiotics can trigger bacterial stress response to trigger dormancy [26]. The transition between adaptive and bet-hedging strategies has been analysed previously [13, 27–29]. It will be interesting to extend the current model (dormancy can be triggered by antibiotic-independent stress or entered spontaneously) to include antibiotic-triggered dormancy.

When we compare the current results with the experimental observations, we should note that the lag time may not be determined purely from growth optimization. For example, *Escherichia coli*’s lag time varies depending on physiological conditions, e.g., the growth medium, rate of environmental changes, and and the duration of the starvation period [30–33]. This indicates that there might be a lower boundary in the lag time for the cell to prepare for regrowth. It has been proposed that spontaneous dormancy may be due to positive feedback loops in the metabolic network, which sometimes drive systems out of growth states. Hence, it may be a side effect of complex metabolic dynamics [34–36]. However, evolutionary experiments have shown that bacteria can prolong their lag time to survive antibiotic application [4, 11]. For spontaneous dormancy, a mutant *hipQ* has been isolated that has a higher subpopulation of spontaneous persisters [37, 38]. There are likely various molecular mechanisms behind cell dormancy, some of which are unavoidable side effects. Therefore, a large bacterial population will always have some level of triggered and spontaneous dormancy. However, if a higher frequency of dormancy is beneficial in a given environment, bacteria can evolve to increase it. Our analysis is more relevant in an environment where high persistence is expected and can help guide an antibiotic administration strategy that avoids selecting for a higher frequency of triggered and/or spontaneous persister cells.

## DATA AVAILABILITY

The datasets generated and analysed during the current study are available in https://github.com/siljabl/BacterialPersistence.

## ACKNOWLEDGMENTS

This research was funded by VILLUM FONDEN (00028054) and the Novo Nordisk Foundation (NNF21OC0068775).

## Appendix A: The analytical solution of two-state model

We rewrite eqs. (1)-(3) to a set of second order differential equations

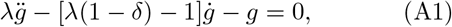

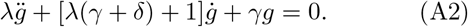

From these we obtain the characteristic equations

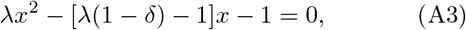

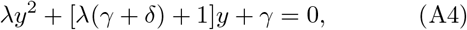

with the solutions

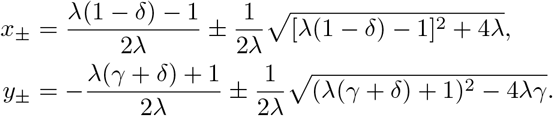

We define the more convenient *a*_*±*_ and *a*_*p±*_
From which we can express the bacterial parameters as

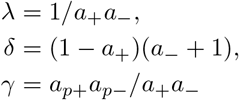

The solutions to eqs. (A1)-(A2) are on the form

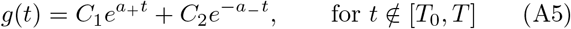

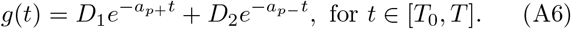

### Solution for t ≤ T_0_

The dynamics before antibiotics are applied to the system follow eq. (A5) with

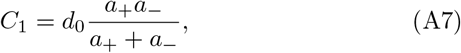

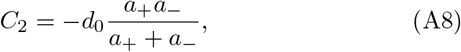

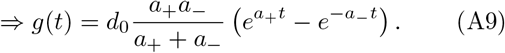

### Solution for t ∈ [T_0_, T]

During the antibiotics the system follows eq. (A6) with

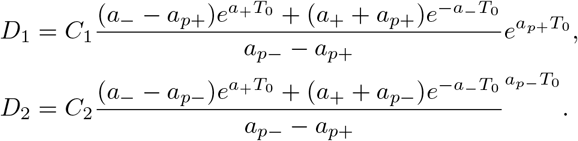

### Solution for t ≥ T

After antibiotics the system again follows eq. (A5), but with

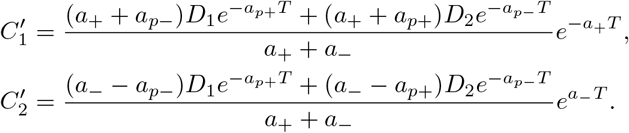

### Computing *T*_*s*_

*T*_*s*_ is the time it takes a single population to consume all the available nutrients, and is found by integrating eq. (3) for *t ∈* [0, *T*_*s*_] and isolate for *T*_*s*_

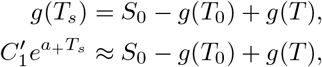

where the second term of *g*(*T*_*s*_) is omitted because 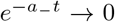 when *t* is large. This approximation is justified in Supplementary Sec. S2. Isolating *T*_*s*_

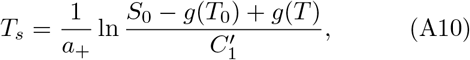

with

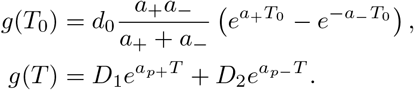

In the absence of antibiotics, eq. (B14) reduces to

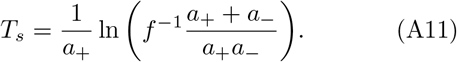

## Appendix B: The analytical solution of three-state model

The three-state model is to a large extent solved in analogue with appendix A, with *ω* replacing *λ* in the second order differential equations (A1)-(A2). The contribution from the triggered persistence turns the system into a set of inhomogeneous second-order differential equations

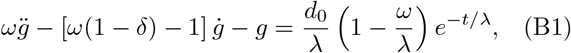

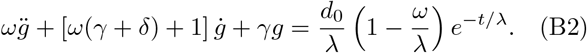

Solving this exactly as in the two-state case, we obtain

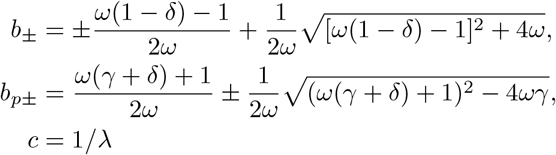

We can express the bacterial parameters as

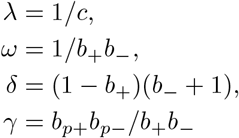

The solutions to eqs. (B1)-(B2) are on the form

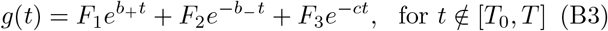

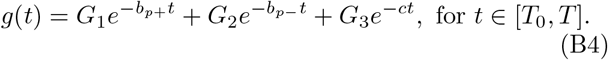

### Solution for t ≤ T_0_

The dynamics before antibiotics are applied to the system follow eq. (B3) with

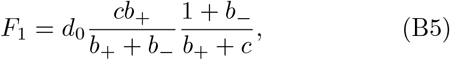

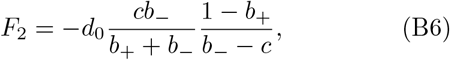

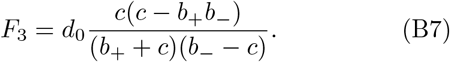

### Solution for t ∈ [T_0_, T]

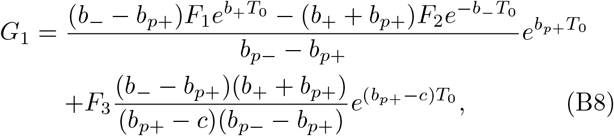

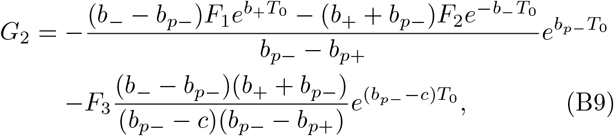

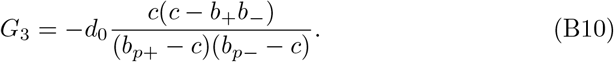

### Solution for t *≥* T

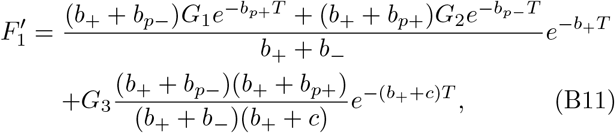

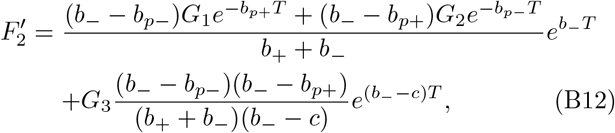

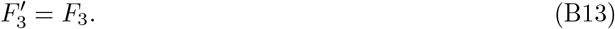

### Computing *T*_*s*_

Like in the two-state model

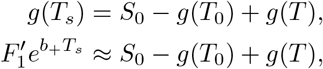

where the second term of *g*(*T*_*s*_) is omitted because 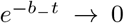 and *e*^−*ct*^ → 0 when *t* is large. These approximations are also justified in Supplementary Sec. S2. Isolating *T*_*s*_

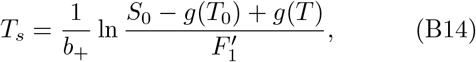

with

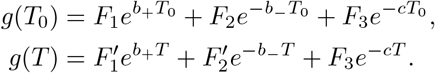

