## Supplementary text for "Triggered and Spontaneous Dormancy in Bacteria During Feast-Famine Cycles with Stochastic Antibiotic Application"

### Supplementary material for: Optimal triggered and spontaneous persistence strategies for bacteria under feast-famine cycles

Silja Borring Låstad and Namiko Mitarai

July 1, 2025

#### Contents


| | | |
|---|---|---|
| S1 | Consumption rate of the nutrient | 2 |
| S2 | Approximations $e^{-const \cdot T_s} \to 0$ | 6 |
| S3 | Fitness of the two-state optimal strategy | 8 |
| S4 | The measure of competition fitness | 9 |
| S5 | The competition optimal strategy | 10 |
| S6 | Two-state mutation | 13 |
| S7 | Fitness of the three-state optimal strategy | 16 |
| S8 | Simulations with stochastic application time | 17 |
| S9 | Mutation rate $\varepsilon = 10^{-2}$ | 19 |
| S10 | Comparing fitness of persistence strategies | 21 |

### S1 Consumption rate of the nutrient

In our model, the rate of nutrient consumption is proportional to changes in the growth state

$$\frac{d}{dt}S(t) = \begin{cases} -\sum_{i=1}^{n} \dot{g}_i(t), & t \notin [T_0, T] \\ 0. & t \in [T_0, T] \end{cases} \tag{1}$$

This implies that bacteria in the growth state consumes nutrients to grow, but also that dormant and persistent bacteria consume nutrients when they undergo phenotypic switching to the growth state. Our model choice relies on the assumption that the rates of phenotypic switching are negligible compared to the growth rate, in which case we obtain

$$\frac{d}{dt}S(t) \approx \begin{cases} -\sum_{i=1}^{n} (1-\delta) g_i(t), & t \notin [T_0, T] \\ 0, & t \in [T_0, T] \end{cases} \tag{2}$$

i.e. a consumption rate that is proportional to the growth rate. Here, we solve the model with consumption rate eq. (2) to confirm that the effect of phenotypic switching is indeed negligible.

#### Two-state model

First, we solve eq. (2) for a single species in the two-state model. Integrating over $t \in [0, T_s]$ and dividing through by $(1-\delta)$ yields

$$\begin{aligned} \frac{S_0}{1-\delta} &= \int_0^{T_0} \left(C_1 e^{a_+ t} + C_2 e^{-a_- t}\right) dt + \int_T^{T_S} \left(C_1' e^{a_+ t} + C_2' e^{-a_- t}\right) dt, \\ \frac{S_0}{1-\delta} &= \left(\frac{C_1}{a_+} e^{a_+ t} - \frac{C_2}{a_-} e^{-a_- t}\right)_0^{T_0} + \left(\frac{C_1'}{a_+} e^{a_+ t} - \frac{C_2'}{a_-} e^{-a_- t}\right)_T^{T_s}, \end{aligned}$$

where $C_i, C_i'$, and $a_\pm$ are as defined in main text Appendix A. Then, we isolate $T_s$ by approximating $e^{-a_- T_s}$ to 0

$$T_s = \frac{1}{a_+} \log\left[\frac{a_+}{C_1'}\left(\frac{S_0}{1-\delta} - \frac{C_1}{a_+}(e^{a_+ T_0} - 1) + \frac{C_2}{a_-}(e^{-a_- T_0} - 1) - \frac{C_2'}{a_-} e^{-a_- T}\right)\right] + T$$

In absence of antibiotics, this simplifies to

$$T_s = \frac{1}{a_+} \log\left(f^{-1} \frac{a_+ + a_-}{a_+ a_-}\right) + \frac{1}{a_+} \log\left(\frac{a_+}{1-\delta} - f\right), \tag{3}$$

where the first term is identical to $T_s$ in the main paper, and the second term is $\approx -f$ when $\delta = 0$.

In Fig. S1 we compare the optimal strategy for the model with consumption rate from eq. (2) with the result in the main text using the consumption rate from eq. (1). The optimal strategy is qualitatively invariant to the choice of

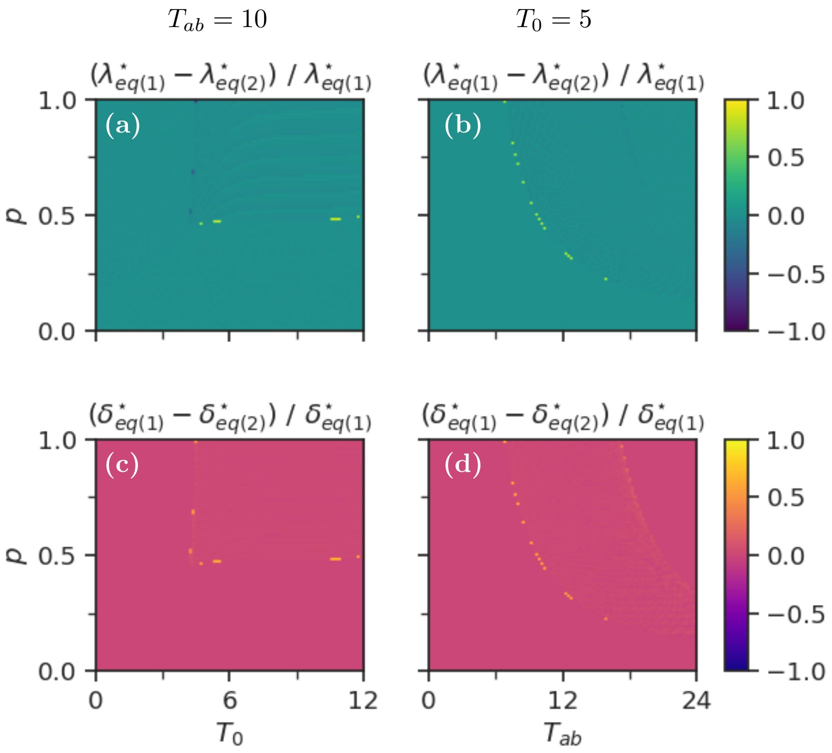


Figure S1 : Comparison of optimal strategies using consumption rate from eq. (1): $\dot{S} = -\sum_i \dot{g}_i$ and from eq. (2): $\dot{S} = -\sum_i (1-\delta_i) g_i$. (a,c) The optimal lag time ($\lambda^\star$), and (b,d) the rate to enter spontaneous dormancy ($\delta^\star$).

consumption rate. The phase boundary shifts slightly, and both $\lambda^\star$ and $\delta^\star$ take slightly different values when modeling the consumption rate as eq. (2). However, these effects are very small, hence for our purposes we conclude that eq. (1) is equivalent to eq. (2).

#### Three-state model

The three-state model is solved in analogue with the two-state model

$$\frac{S_0}{1-\delta} = \int_0^{T_0} (F_1 e^{b_+ t} + F_2 e^{-b_- t} + F_3 e^{-ct}) dt + \int_T^{T_s} (F_1' e^{b_+ t} + F_2' e^{-b_- t} + F_3 e^{-ct}) dt, \tag{4}$$

where $F_i, F_i', b_\pm$, and $c$ are as defined in main text Appendix B. We isolate $T_s$ by approximating $e^{-b_- T_s}$ and $e^{-c-T_s}$ to 0.

$$T_s = \frac{1}{b_+} \log \left[ \frac{b_+}{F_1'} \left( \frac{S_0}{1-\delta} - \frac{F_1}{b_+}(e^{b_+ T_0} - 1) + \frac{F_2}{b_-}(e^{-b_- T_0} - 1) + \frac{F_3}{c}(e^{-cT_0} - e^{-cT} - 1) - \frac{F_2'}{b_-} e^{-b_- T} \right) \right] + T. \tag{5}$$

Finally, we compare the optimal strategy for the model with consumption rate eq. (2) with the result in the main text using consumption rate eq. (1). The result in shown in Fig. S2 , and we confirm that also for the three-state model is the effect from including phenotypic switching negligible.

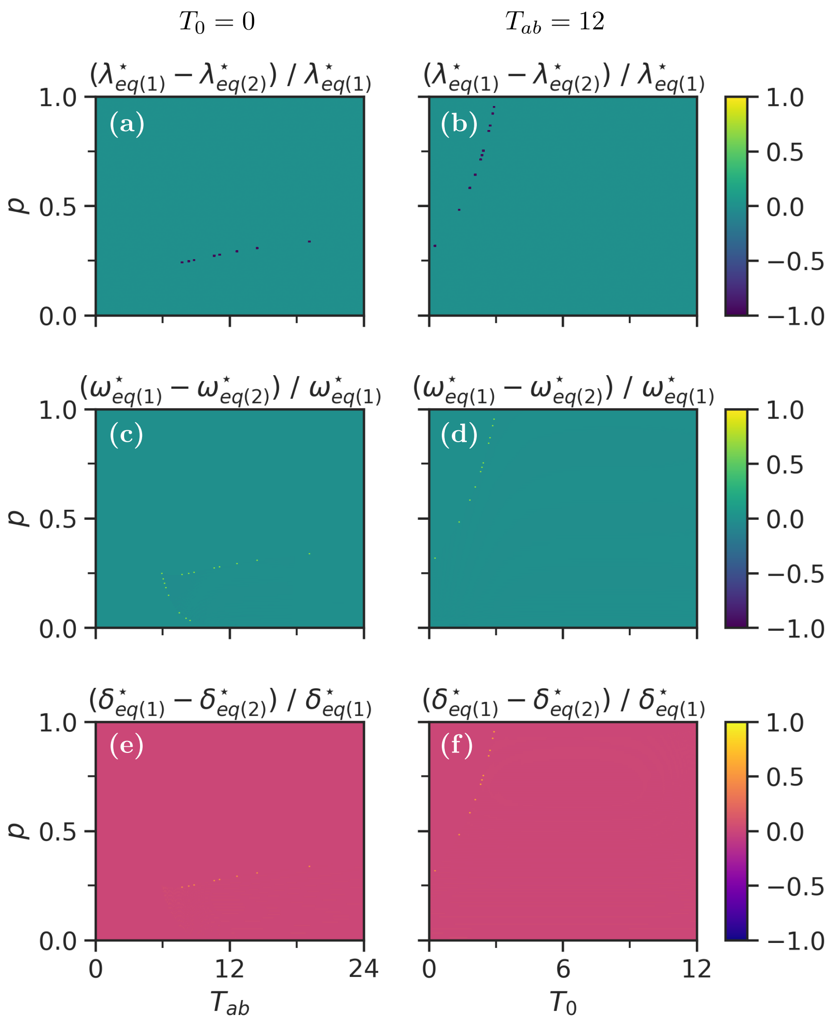


Figure S2 : Comparison of optimal strategies using consumption rate from eq. (1): $\dot{S} = -\sum_i \dot{g}_i$ and from eq. (2): $\dot{S} = -\sum_i (1-\delta_i) g_i$. (a,d) The lag time ($\lambda^\star$), (b,e) the time spent in spontaneous dormancy ($\omega^\star$), and (c,f) the rate to enter the spontaneous dormancy ($\delta^\star$).

### S2 Approximations $e^{-const \cdot T_s} \to 0$

Here we justify the approximations $e^{-a_- T_s} \to 0$, $e^{-b_- T_s} \to 0$, and $e^{-cT_s} \to 0$ that are used to compute the single species optimal strategy in the two- and three-state models, respectively.

#### Two-state model

##### $\mathbf{e^{-a_- T_s} \to 0}$

We solve the two-state model by approximating

$$g(T_s) = C_1' e^{a_+ T_s} + C_2' e^{-a_- T_s} \approx C_1' e^{a_+ T_s}, \tag{6}$$

where the constants $C_i'$ and $a_\pm$ are given in main text Appendix A.

To see that $e^{-a_- T_s} \to 0$, we study the ratio of the two exponentials

$$\frac{e^{-a_- T_s}}{e^{a_+ T_s}} \leq \left( \exp\left[ -\frac{\sqrt{1+4T}}{T} \right] \right)^{T_s} \approx \left( \exp\left[ -2/\sqrt{T} \right] \right)^{T_s}, \tag{7}$$

i.e. the ratio goes exponentially to 0. The upper boundary on the ratio corresponds to the case with $\delta = 1$ and $\lambda = T$, where we have assumed that $T$ is the largest realistic value that $\lambda$ can take because in our model there is no further advantage from having $\lambda > T$. However, we can only neglect the term with $e^{-a_- T_s}$ if the factor in front is sufficiently small. Comparing $C_2'$ with $C_1'$, we see that $C_2' < C_1'$ for all parameters

$$\left| \frac{C_2' e^{-a_- T}}{C_1' e^{a_+ T}} \right| = \left| \frac{(a_- - a_{p-}) D_1 e^{-a_{p+} T} + (a_- - a_{p+}) D_2 e^{-a_{p-} T}}{(a_+ + a_{p-}) D_1 e^{-a_{p+} T} + (a_+ + a_{p+}) D_2 e^{-a_{p-} T}} \right| < 1.$$

We conclude that terms on the form $e^{-a_- T_s}$ can be neglected.

#### Three-state model

##### $\mathbf{e^{-b_- T_s} \to 0}$

We do the equivalent approximation in the three-state model

$$g(T_s) = F_1' e^{b_+ T_s} + F_2' e^{-b_- T_s} + F_3' e^{-cT_s} \approx F_1' e^{b_+ T_s}, \tag{8}$$

where the constants $F_i', b_\pm$ and $c$ are given in man text Appendix B.

$e^{-b_- T_s}/e^{b_+ T_s}$ follow eq. 7, therefore we can ignore the $e^{-b_- T_s}$-term if the ratio $F_2'/F_1'$ does not diverge. From the ratio

$$\frac{F_2' e^{-b_- T}}{F_1' e^{b_+ T}} = \frac{(b_- - b_{p-}) G_1 e^{-b_{p+} T} + (b_- - b_{p+}) G_2 e^{-b_{p-} T} + G_3 \frac{(b_- - b_{p-})(b_- - b_{p+})}{b_- - c} e^{-cT}}{(b_+ + b_{p-}) G_1 e^{-b_{p+} T} + (b_+ + b_{p+}) G_2 e^{-b_{p-} T} + G_3 \frac{(b_+ + b_{p-})(b_+ + b_{p+})}{b_+ + c} e^{-cT}},$$

we see that the ratio blows up when $b_- = c$, but is otherwise smaller than 1. We express $c$ as $c = b_- - \epsilon$, where $\epsilon \ll 1$, and treat this case together with the $e^{-cT_s}$-term in the next section.

$\mathbf{e^{-cT_s} \to 0}$

For any combination of bacterial parameters we have

$$\frac{e^{-cT_s}}{e^{b_+T_s}} \leq \left(\exp\left[-\frac{1+\sqrt{1+4T}}{2T}\right]\right)^{T_s} < \left(\exp\left[-1/\sqrt{T}\right]\right)^{T_s}, \tag{9}$$

which also goes exponentially to 0.

Then, we study the ratio of of factors in front of the exponentials, comparing only the $c$-dependent term of $F_1'$ with $F_3$ for simplicity.

$$\left|\frac{F_3}{F_1' e^{b_+T}}\right| < \frac{G_3 \frac{(b_{p+}-c)(b_{p-}-c)}{(b_++c)(b_--c)}}{G_3 \frac{(b_++b_{p-})(b_++b_{p+})}{(b_++b_-)(b_++c)}} = \frac{(b_++b_-)(b_{p+}-c)(b_{p-}-c)}{(b_--c)(b_++b_{p-})(b_++b_{p+})},$$

where $F_3$ has been expressed as a function of $G_3$ and the strict inequality comes from the two additional terms of $F_1'$ that are not considered here. The ratio blows up when $c \to \infty$ and $b_- = c$, but is otherwise smaller than 1. We can disregard the case $c \to \infty$ because the exponential $e^{-cT_s}$ goes faster to 0.

Finally, we consider the case $c = b_- - \epsilon$, $\epsilon \ll 1$, and study the two terms that diverge in this region, namely $F_2' e^{-b_-t}$ and $F_3 e^{-ct}$. $F_2'$ has three terms, but we focus on the last term in main text eq. (B12) because the two other are independent of $c$. Adding the last term of $F_2'$ to $F_3$, and substituting $c = b_- - \epsilon$, yields

$$= \frac{G_3}{\epsilon}\left(\frac{(b_- - b_{p-})(b_- - b_{p+})}{b_+ + b_-} e^{\epsilon T} - \frac{(b_{p+} - b_- + \epsilon)(b_{p-} - b_- + \epsilon)}{b_+ + b_- - \epsilon} e^{\epsilon T_s}\right) e^{-b_-T_s}.$$

We expand this around $\epsilon = 0$

$$= \frac{G_3}{\epsilon}\left(\frac{(b_- - b_{p-})(b_- - b_{p+})}{b_+ + b_-}\epsilon(T - T_s) - \frac{(b_{p+} + b_{p-} - 2b_-)}{b_+ + b_-}\epsilon - \frac{(b_- - b_{p-})(b_- - b_{p+})}{(b_+ + b_-)^2}\epsilon\right) e^{-b_-T_s},$$

$$= -\frac{G_3}{b_+ + b_-}\left((b_- - b_{p-})(b_- - b_{p+})\left(T_s - T + \frac{1}{b_+ + b_-}\right) + b_{p+} + b_{p-} - 2b_-\right),$$

which is well behaved and reasonably small for all meaningful parameters.

#### S3 Fitness of the two-state optimal strategy

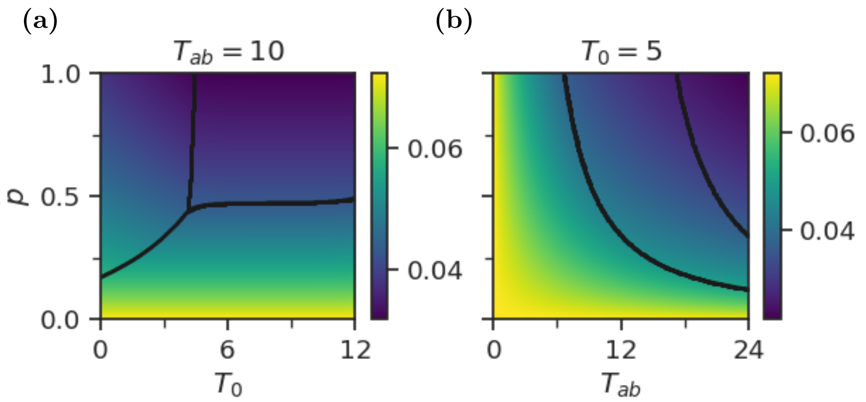


Figure S3 : The fitness corresponding to the optimal persistence strategies for the three-state model. The black lines illustrate the phase boundary. a) Varying $p$ and $T_0$, while keeping $T_{ab} = 10$. b) Varying $p$ and $T_{ab}$, while keeping $T_0 = 5$.

The fitness corresponding to the optimal persistence strategies for the antibiotic parameters explored in the main text is shown in Fig. S3 . Fitness decreases with increasing antibiotic severeness, as expected. The decrease is continuous, hence, phase transitions occur when the fitness of one strategy overtakes the fitness of another in a continuous manner. Varying $T_0$ has little effect on the fitness.

### S4 The measure of competition fitness

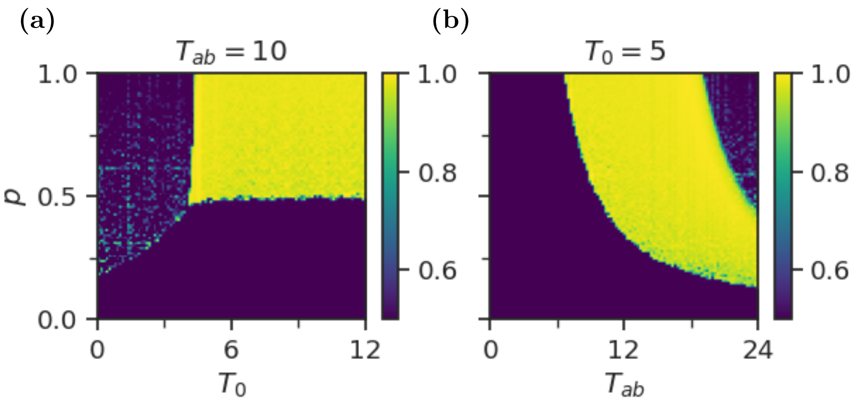


Figure S4 : Fraction of nutrients that is consumed by the competitor competing against the single species optimal persistence strategy for a) $T_{ab} = 10$, and b) $T_0 = 5$.

The cycle average of how much nutrients were converted to the biomass of the best competitor species is plotted in Fig. S4 . The value ranges between 0.5 and 1, where 0.5 implies that the optimal strategy for a single species is also the optimal strategy for competition (the single species optimum and the competitor species are identical and thus consume exactly the same amount of nutrients).

In Fig. S4 we observe three phases in the fraction of converted biomass corresponding to the three phases of the optimal dormancy strategy in the main text. Where the single species optimal strategy is to have no dormancy the consumption fraction is $\sim 0.5$, implying that this is also the competition optimal strategy. In the region where the single species optimal strategy is to have only triggered dormancy this is also mostly the case. The fluctuations in consumption fraction here are interpreted as a reflection of the stochastic nature of the system, with $N = 10^4$ cycles not being enough cycles for the winner to dominate the competition fully. However, in the region of both triggered and spontaneous dormancy, the best competitor consumes close to all the nutrients, hence here the competition optimal is different from the single species optimal strategy.

#### S5 The competition optimal strategy

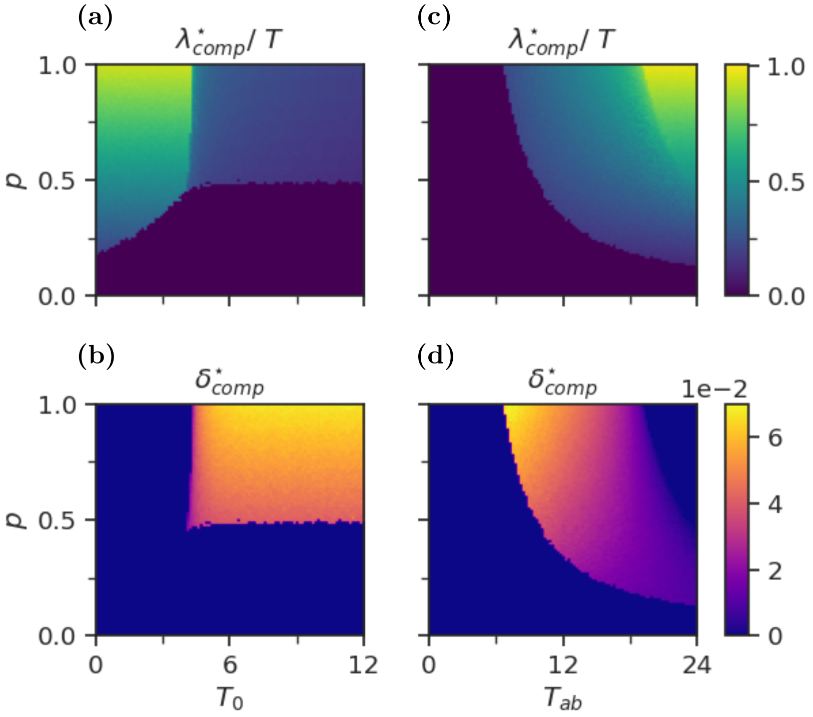


Figure S5 : The optimal persistence parameters for competition in the two-state model for $T_{ab} = 10$ with varying $T_0$ and $p$ (a-b) and with $T_0 = 5$ and varying $T_{ab}$ and $p$ (c-d). (a,c) The optimal lag time ($\lambda^\star_{\text{comp}}$) normalized by $T$, and (b,d) the rate to enter spontaneous dormancy ($\delta^\star_{\text{comp}}$).

In Fig. S5 we plot heatmaps of the optimal persistence strategy when two species compete with each other. The antibiotic parameters are the same as in Fig. 2 in the main text. The competition optimal persistence strategies are qualitatively similar to the single species optimal, but the phase boundary between both spontaneous and triggered dormancy and only triggered dormancy in Fig. S5 b is shifted. This transition occurs at higher $T_{ab}$ when there is competition for nutrients. The corresponding boundary in Fig. S5 a-b ($T_{ab} = 10$) is approximately the same with and without competition. This is observed more easily in Fig. S6 , where we compare the single species optimal persistence parameters with the competition optimal parameters as functions of $T_0$ (a,b) and $T_{ab}$ (c,d). Here, we also observe the increase in $\delta^\star_{\text{comp}}$ compared to $\delta^\star$. The effect of competition on the optimal persistence strategy is qualitatively similar to the effect of increasing the dilution fraction. This is illustrated in Fig. S7 , where the competition optimal strategies from Fig 3 in the main text is plotted against

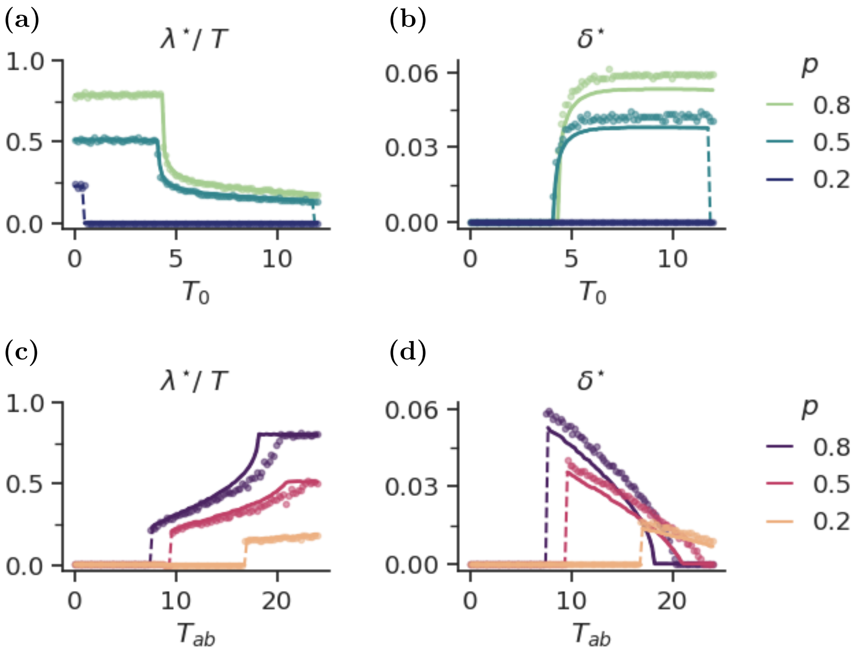


Figure S6 : Optimal lag time and optimal rate of spontaneous dormancy obtained as a function of the antibiotic application time $T_0$ by direct competition of strategies (a-b) or the duration of the antibiotic application (c-d). (a) optimal lag time $\lambda^\star$ and (b) optimal rate of spontaneous dormancy $\delta^\star$ for $T_{ab} = 10$. (c) optimal lag time $\lambda^\star$ and (d) optimal rate of spontaneous dormancy $\delta^\star$ for $T_0 = 5$. The circles denote the competition optimal ($\lambda^\star_{\text{comp}}$ and $\delta^\star_{\text{comp}}$) and the solid lines denote the single species optimal from $F$ (eq. 7).

the single species optimal strategy with dilution fraction $f' = 4f$. Though not equal, the overlap is better than in main text Fig. 3. Adding competition will generally reduce the amount of nutrients available per species. Increasing the dilution fraction will reduce the amount of nutrients compared to the bacterial population at the beginning of a cycle and therefore generally decrease $T_s$ (eq. 4), therefore it seems reasonable that their effect on the optimal strategy is similar.

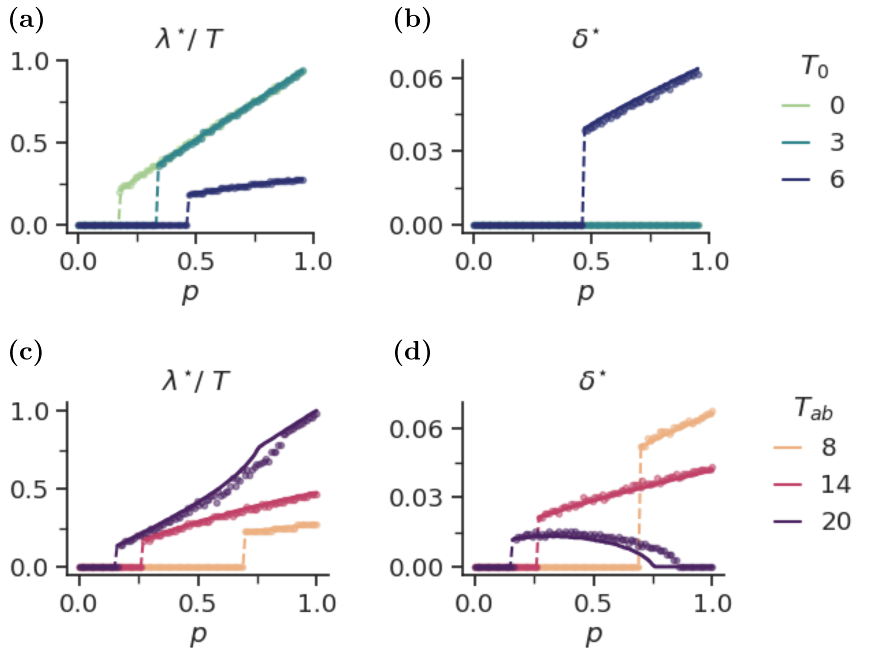


Figure S7 : Competition optimal strategies with dilution fraction $f$ plotted against the single species optimal strategies with dilution fraction $f' = 4f$. The competition optimal strategies are the same as in main text Fig. 3. (a) optimal lag time $\lambda^\star$ and (b) optimal rate of spontaneous dormancy $\delta^\star$ for $T_{ab} = 10$. (c) optimal lag time $\lambda^\star$ and (d) optimal rate of spontaneous dormancy $\delta^\star$ for $T_0 = 5$. The circles denote the competition optimal ($\lambda^\star_{\text{comp}}$ and $\delta^\star_{\text{comp}}$) and the solid lines denote the single species optimal from $F$ (eq. 7).

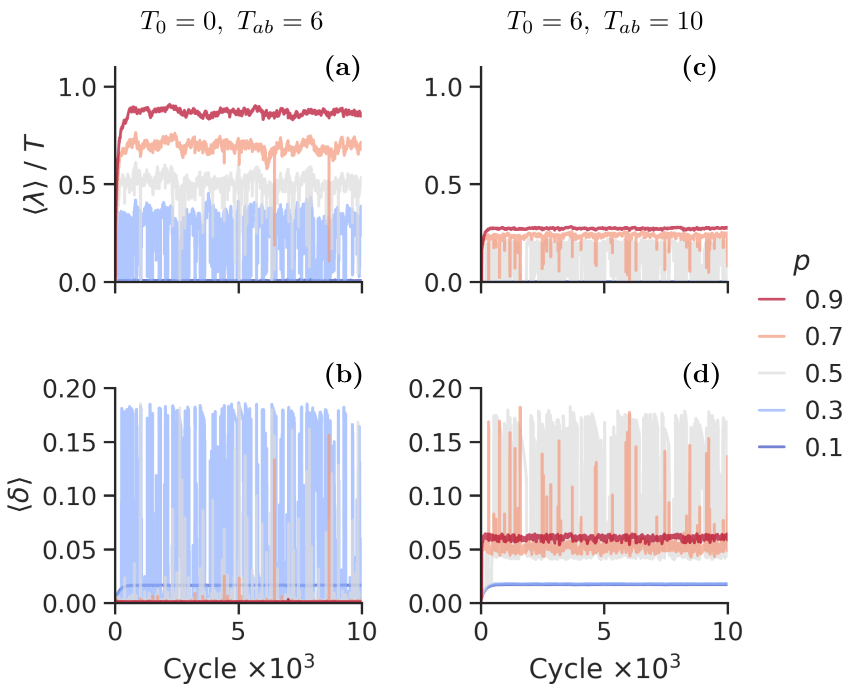


Figure S8 : Evolution of average persistence strategy for $T_0 = 0$ and $T_{ab} = 6$ (a-b) and $T_0 = 6$ and $T_{ab} = 10$ (c-d) with varying $p$ and the mutation rate $\varepsilon = 10^{-3}$. (a,c) The average lag time ($\langle\lambda\rangle$) normalized by $T$, (b,d) the rate to enter the spontaneous dormancy ($\langle\delta\rangle$). Different lines correspond to different values of $p$, as labelled to the figure's right.

### S6 Two-state mutation

We consider mutation in the two-state model with the same setup as in main text Fig. 6. We let a population to have 2 indices $(i, j)$, which has a parameter set

$$(\lambda_i, \delta_j) = (i\Delta + 0.01, j\Delta_\delta) \tag{10}$$

with $\Delta = 0.2$ and $\Delta_\delta = 0.01$. Then, in the absence of antibiotics, the growing part of the population $(i, j)$ obeys

$$\frac{d}{dt}g_{i,j}(t) = \frac{d_{i,j}(t)}{\lambda_i} + (1 - \delta_j - \eta\varepsilon)g_{i,j}(t) + \varepsilon \sum_l \left[g_{i+l,j}(t) + g_{i,j+l}\right]. \tag{11}$$

Four sets of parameters are explored in Fig. S8 -S9 . In Fig. S8 a-b we consider the case of $T_0 = 0$ and $T_{ab} = 6$. For $p \geq 0.5$ we obtain only triggered dormancy, though with some fluctuations. For $p = 0.1$ the optimal strategy is no dormancy, and for $p = 0.3$ the system is constantly fluctuating between no dormancy and triggered dormancy.

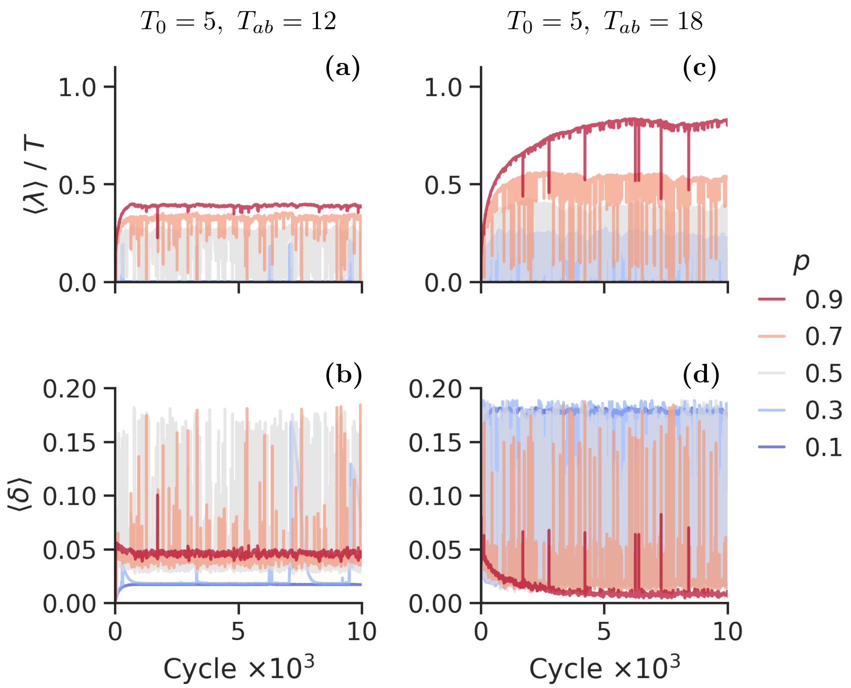


Figure S9 : Evolution of average persistence strategy for $T_0 = 5$ and $T_{ab} = 12$ (a-b) and $T_0 = 5$ and $T_{ab} = 18$ (c-d) with varying $p$ and the mutation rate $\varepsilon = 10^{-3}$. (a,c) The average lag time ($\langle\lambda\rangle$) normalized by $T$, (b,d) the rate to enter the spontaneous dormancy ($\langle\delta\rangle$). Different lines correspond to different values of $p$, as labelled to the figure's right.

In Fig. S8 c-d we plot $T_0 = 6$ and $T_{ab} = 10$. This corresponds to antibiotic parameters along a vertical line in main text Fig. 2a-b, from which we expect no dormancy for $p < 0.5$ and both triggered and spontaneous dormancy for $p > 0.5$. This is also what we observe in Fig. S8 c-d. For $p = 0.5$ the system is heavily fluctuating between the two optimal strategies. From main text Fig. 2a-b this is not unexpected, as $p = 0.5$ is very near the phase boundary. Like in the three-state model $\langle\delta\rangle$ is increasing when $\langle\lambda\rangle$ is decreasing.

In Fig. S9 we investigate parameter combinations from main text Fig. 2c-d. In Fig. S9 a-b we plot the case of $T_0 = 5$ and $T_{ab} = 12$, for which we expect both triggered and spontaneous dormancy for $p \geq 0.5$, no dormancy for $p \leq 0.3$. This is mostly what we observe in Fig. S9 a-b. For $p = 0.5$ the system is again heavily fluctuating.

In Fig. S9 c-d we consider $T_0 = 5$ and $T_{ab} = 18$. From the analytical solution in main text Fig. 2c-d we expect only triggered dormancy for $p \geq 0.7$, both triggered and spontaneous for $p = 0.3 - 0.5$ and no dormancy for $p = 0.1$. For $p = 0.9$ the mutation simulation does not reach a steady state within $10^4$ fest-famine cycles. $\langle\delta\rangle$ is decreasing towards a small but finite value, whereas $\langle\lambda\rangle$ is slowly increasing towards a value between the triggered optimal ($\lambda \sim pT$) and the mixed strategy optimal value. $p = 0.9$ is close to the phase boundary between only triggered and mixed persistence strategy. From Section S5 we know that competition can move this boundary, which might explain the deviation from the analytical optimal strategy observed here.

For $p = 0.3 - 0.7$ the system is heavily fluctuating around its mixed persistence optimal values. This is similar to what we observe in Fig. S9 a-b, but both the amplitude and frequency are higher for $T_{ab} = 18$. Finally, for $p = 0.1$ we observe $\langle\lambda\rangle \to 0$, but $\langle\delta\rangle \approx 0.18$. Hence, for rare, but severe antibiotic events there seems to be a competitive advantage of having a finite rate to enter spontaneous dormancy, though this is not explored further here.

#### S7 Fitness of the three-state optimal strategy

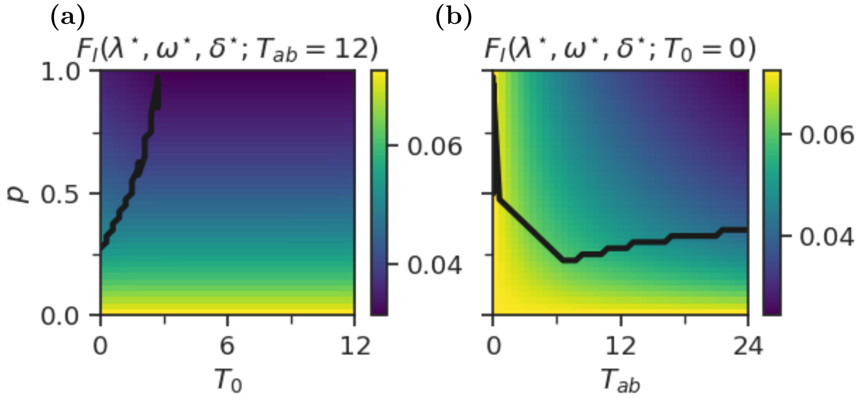


Figure S10 : The fitness corresponding to the optimal persistence strategies for three-state model. The black lines illustrate the phase boundary.

Fitness heatmaps corresponding to the optimal persistence strategies in the three-state model are shown in Fig. S10 . The heat maps are similar to Fig. S3 in that fitness continuously decreases as the severity of the antibiotic increases. Again, $T_0$ does not appear to have an effect on the optimal fitness.

#### S8 Simulations with stochastic application time

We run simulations where we allow $T_0$ to fluctuate between two values: $T_0 = 0$, representing the case of synchronized addition of antibiotics and nutrients, and $T_0 > 0$, representing the desynchronized case of delayed antibiotics. Antibiotics are delayed with probability $p_{T_0}$ and are otherwise synchronized. We run the simulation for $p_{T_0} \in [0, 1]$ for various $T_0$ and $p$ and study the optimal persistence strategy.

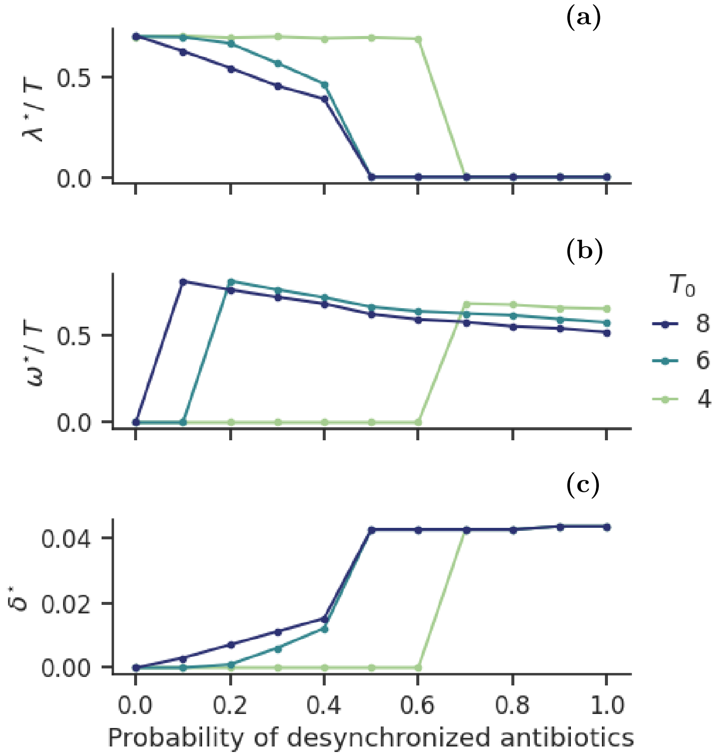


Figure S11 : Optimal persistence parameters when the application time of antibiotics $T_0$ is stochastic. $T_{ab} = 12$ and $p = 0.7$. $T_0$ takes the value in the legend when desynchronized, and is 0 otherwise. (a) optimal wake-up rate from triggered dormancy, (b) optimal wake-up rate from spontaneous dormancy, (c) optimal rate to enter spontaneous dormancy.

In Fig. S11 the result is shown for $p = 0.7$, $T_{ab} = 12$ and delayed $T_0 \in \{4, 6, 8\}$. We observe that for $T_0 = 4$ we obtain the same phases as in the case of fixed $T_0$, i.e. only triggered dormancy for $p_{T_0} \leq 0.6$ and otherwise only spontaneous dormancy. For $T_0 = 6$ or 8 the optimal strategy is to have only spontaneous dormancy for $p_{T_0} \geq 0.5$, but below this value we observe a new phase in the three-state model, namely the mixed phase of both triggered and spontaneous dormancy. Here, we have $\lambda^\star > 0$ and $\delta^\star > 0$, but both are smaller than their optimal values in the case of fixed $T_0$. $\omega^\star$ is approximately the same as in the fixed case. Hence, instead of sleeping through the entire antibiotic application, the population can benefit from waking up a little earlier if it has a small rate of entering spontaneous dormancy.

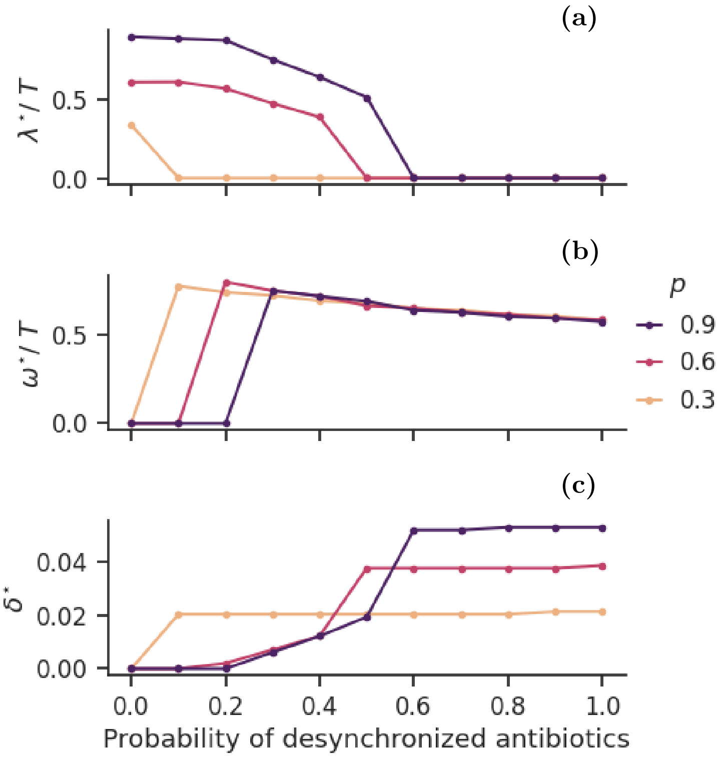


Figure S12 : Optimal persistence parameters when the application time of antibiotics $T_0$ is stochastic. $T_{ab} = 12$, and $T_0 = 6$ when desynchronized and is 0 otherwise. (a) optimal wake-up rate from triggered dormancy, (b) optimal wake-up rate from spontaneous dormancy, (c) optimal rate to enter spontaneous dormancy.

Fig. S12 shows the results from the simulation with $T_0 = 6$, $T_{ab} = 12$ and $p \in \{0.3, 0.6, 0.9\}$. For $p = 0.3$ we observe only spontaneous dormancy for all $p_{T_0} > 0$. For both $p = 0.6$ and 0.9 there is a range of $p_{T_0}$ that yields a mixed strategy as the best strategy. As in Fig. S11 , $\lambda^\star$ and $\delta^\star$ take values that are lower than the optimal in the fixed case, while $\omega^\star$ is approximately equal to the purely spontaneous optimal.

#### S9 Mutation rate $\varepsilon = 10^{-2}$

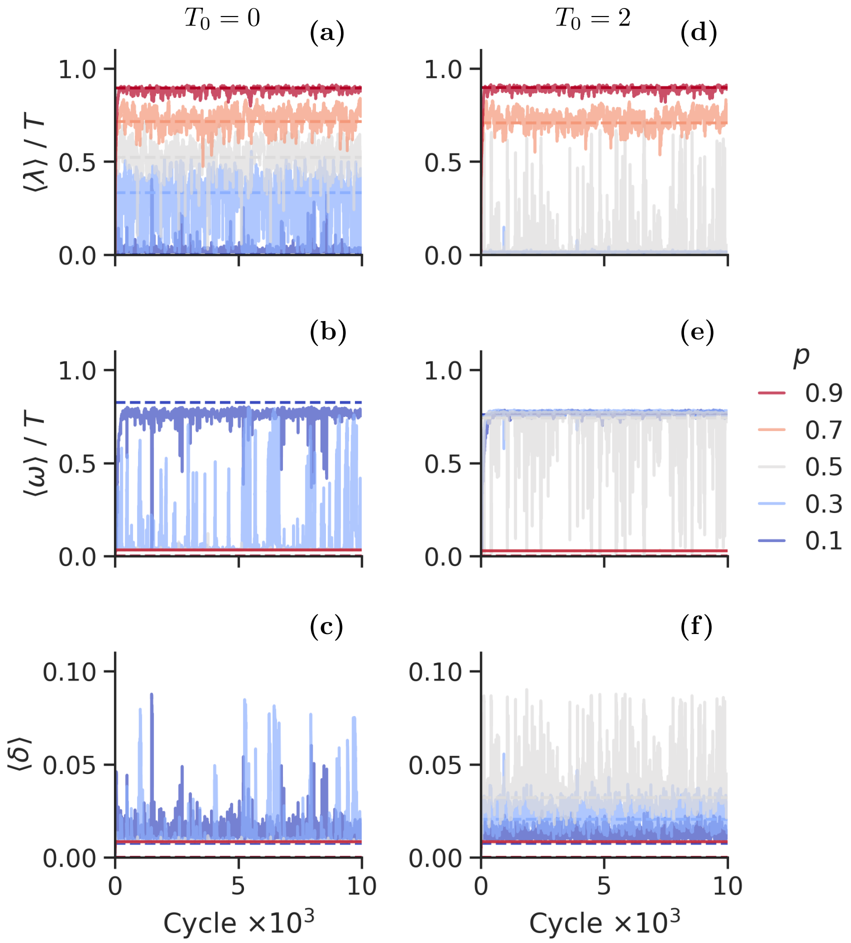


Figure S13 : Evolution of average persistence strategy for $T_0 = 0$ and $T_{ab} = 12$ (a-c) and $T_0 = 2$ and $T_{ab} = 12$ (d-e) with varying $p$ and the mutation rate $\varepsilon = 10^{-3}$. (a,d) The average lag time ($\langle\lambda\rangle$) normalized by $T$, (b,e) the time spent spontaneous dormancy ($\langle\omega\rangle$) normalized by $T$, and (c,f) the rate to enter the spontaneous dormancy ($\langle\delta\rangle$). Different lines correspond to different values of $p$, as labelled to the figure's right. The set of antibiotic parameters explored are highlighted as coloured circles in main text Fig. 5.

Here, we explore the effect of increasing the mutation rate to $\varepsilon = 10^{-2}$, which is in the range of $\delta^\star$. The result is shown in Fig. S13 . With higher mutation rate, the fluctuations around the optimal values become more frequent and with larger amplitude for most parameter sets. Fig. S13 mostly overlap with the single species optima in main text Fig. 5. The exception is $T_0 = 0, p = 0.1$, where $\langle\omega\rangle < \omega^{\star}$. Increasing the mutation rate increases the competition from other species, hence it seems that competition may alter the optimal strategy slightly also in the three-state model, similar to what we observed in the two-state model.

#### S10 Comparing fitness of persistence strategies

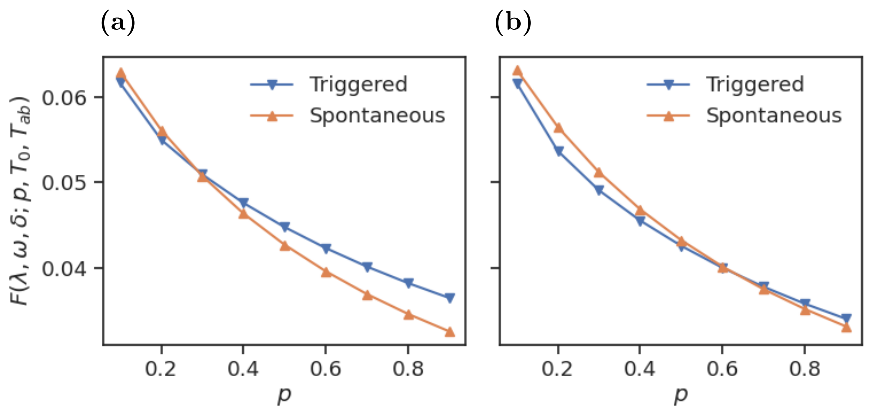


Figure S14 : The fitness of the best triggered and spontaneous strategies, respectively. (a) $T_0 = 0$, $T_{ab} = 12$, (b) $T_0 = 2$, $T_{ab} = 12$

In Fig. S14 we compare the fitness of the best triggered strategy with the fitness of the best spontaneous strategy for several $p$. We observe that both are decreasing as the frequency of antibiotics increases, but that the fitness of the purely spontaneous persistence strategy is decreasing faster than that of the purely triggered persistence strategy.

In Figs. S15 -S16 we study the fitness dependence on $\lambda$ and $\omega$. In order to facilitate visualization, we have made a maximum projection of the fitness along the $\delta$-axis. The corresponding maximal $\delta$ is plotted along with the fitness. The best triggered persistence strategy is highlighted by the cyan arrow on the $x$-axis, and the best spontaneous strategy is highlighted by the orange arrow on the $y$-axis. The fitness landscape varies continuously, and takes approximately identical values for several $\lambda$ and $\omega$. As expected, varying $\omega$ has no effect on the fitness where $\delta = 0$. Also varying $\lambda$ has little effect on the fitness, but as seen in the two-state competition simulation, even an infinitesimal advantage will matter when the number of feast-famine cycles becomes large. However, this implies that in a more realistic scenario in which also extinction is considered the trade-off of giving up a little fitness in order to be better protected against extinction might indeed be beneficial.

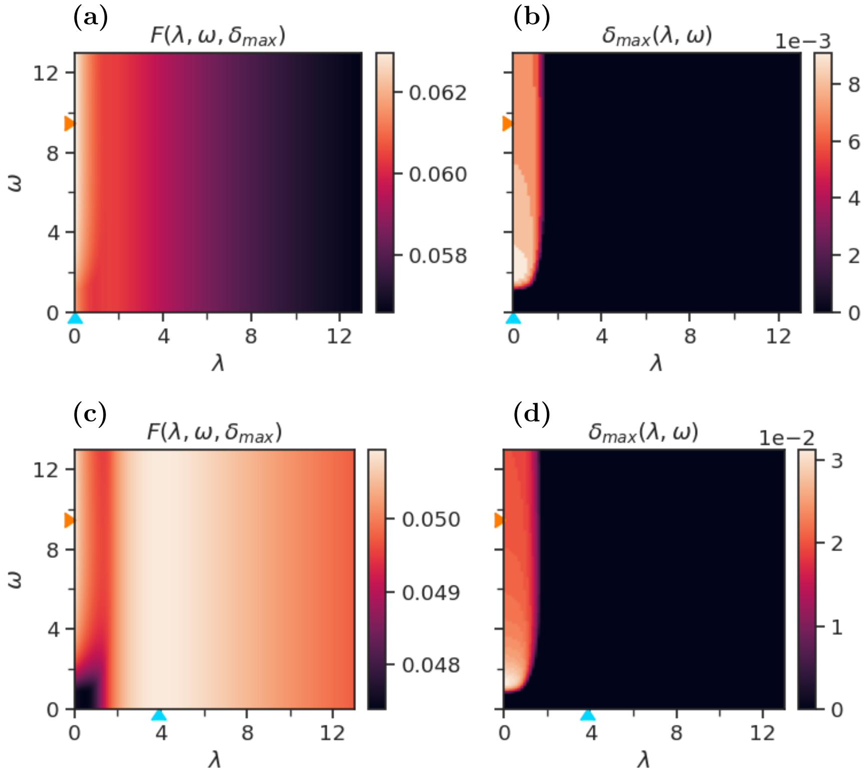


Figure S15 : Fitness and $\delta$ heatmaps as function of $\lambda$ and $\delta$. A maximal projection is taken on the 3-dimensional fitness matrix along the $\delta$-axis to reduce it to 2-dimensional. The corresponding $\delta_{\max}$ is plotted on the right. Orange arrows represent optimal spontaneous lag time $\omega$, and cyan arrows represent optimal triggered lag time $\lambda$. Here, $T_0 = 0$ and $T_{ab} = 12$. (Top) $p = 0.1$. (Bottom) $p = 0.3$

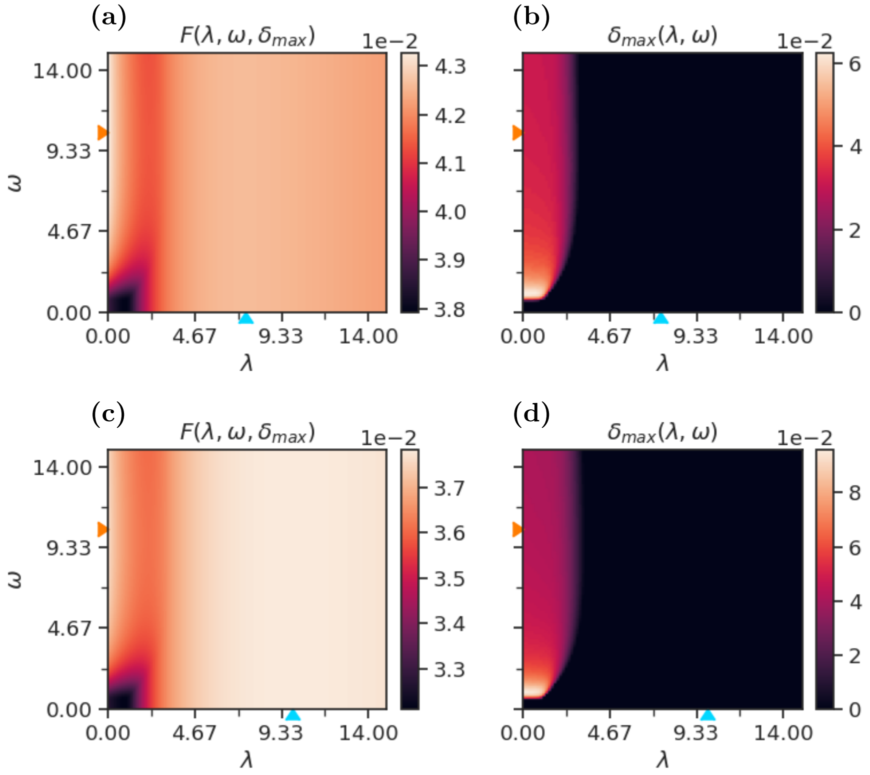


Figure S16 : Fitness and $\delta$ heatmaps as function of $\lambda$ and $\delta$. A maximal projection is taken on the 3-dimensional fitness matrix along the $\delta$-axis to reduce it to 2-dimensional. The corresponding $\delta_{\max}$ is plotted on the right. Orange arrows represent optimal spontaneous lag time $\omega$, and cyan arrows represent optimal triggered lag time $\lambda$. Here, $T_0 = 2$ and $T_{ab} = 12$. (Top) $p = 0.5$. (Bottom) $p = 0.7$
